# The intrinsic dimension of gene expression during cell differentiation

**DOI:** 10.1101/2024.08.02.606382

**Authors:** Marta Biondo, Niccolò Cirone, Filippo Valle, Silvia Lazzardi, Michele Caselle, Matteo Osella

## Abstract

Waddington’s epigenetic landscape has long served as a conceptual framework for understanding cell fate decisions. The landscape’s geometry encodes the molecular mechanisms that guide the gene expression profiles of uncommitted cells toward terminally differentiated cell types. In this study, we demonstrate that applying the concept of intrinsic dimension to single-cell transcriptomic data can effectively capture trends in expression trajectories, supporting this framework. This approach allows us to define a robust cell potency score without relying on prior biological information. By analyzing an extensive collection of datasets from various species, experimental protocols, and differentiation processes, we validate our method and successfully reproduce established hierarchies of cell type potency.

## Introduction

The Waddington’s epigenetic landscape is often used as a metaphor to rationalize the cell differentiation process [1]. Uncommitted progenitor cells progressively lose their potency as they roll down on a high-dimensional surface whose geometry reflects the biological constraints and complex regulations that canalize the differentiation trajectories. Finally, the cells end up in stable basins representing terminally differentiated cell types. Since a cell state is essentially defined by its gene expression profile, expression levels are the natural coordinates in this large epigenetic space [2]. These coordinates, or at least a proxy, can be experimentally measured thanks to recent innovations in single-cell RNA-sequencing (scRNA-seq) [3, 4].

While Waddington’s evocative description has been instrumental in interpreting several results in developmental biology [5], the extent to which this picture is supported by large-scale empirical data remains to be tested. The trajectories followed by single cells in the expression space during differentiation should reflect the presence of an underlying landscape and its specific geometry. However, how to confirm this intuition using RNA sequencing data, how the information about the landscape geometry can be extracted, and how it is related to the differentiation potential are still largely open questions.

Rugged and high-dimensional landscapes, as the one depicted in Waddington’s drawings, are ubiquitous in statistical physics. They typically represent energy surfaces of complex systems such as spin glasses [6, 7], but analogous descriptions are used for fitness landscapes in evolutionary biology [8] or for loss functions of artificial neural networks in machine learning [9]. We plan to leverage on these analogies to identify geometry-based observables that can reveal the presence of an underlying landscape guiding differentiation from scRNA-seq data. Cell positions on the landscape can then be used to define a potency score only relying on data geometry, without the need for prior biological knowledge or well defined “marker genes”, whose expression is often used to annotate cell types [10].

More specifically, the analogy with statistical physics systems suggests that we can consider pluripotent cells as cells in a “high-temperature” state. In fact, they should be able to freely navigate the landscape, only to become progressively constrained into a specific valley when they commit to a differentiation path. Differentiation would correspond to a “freezing process”, during which the expression profiles are narrowed in the manifolds of cell types. This picture agrees with the observation that multipotent cells do not typically have a specific and conserved expression profile or clear-cut markers [11]. Stemness seems rather characterized by pervasive transcription [12] and by a general high level of heterogeneity [13, 14].

This paper will test the presence of a trend that captures the progressive reduction of the accessible expression space during differentiation using the concept of intrinsic dimension. Data associated with complex systems are often high-dimensional, yet statistical regularities stemming from dependencies and correlations tend to concentrate data points on low-dimensional manifolds [15]. Also gene expression data, and particularly RNA-sequencing data, exhibit various statistical laws and correlation patterns [16, 17], suggesting that they can be efficiently described by a number of variables significantly smaller than the large gene repertoire. This notion has been previously suggested [18, 19], and it is implicitly assumed by most data preprocessing pipelines that incorporate dimensionality reduction steps [20].

The hypothesis this paper focuses on is that the intrinsic dimension of cell expression profiles decreases with cell differentiation, and thus this quantity can be used to robustly measure the potency of a cell population. Drawing an analogy between the Waddington’s landscape and the energy profile of statistical physics models, such as the Hopfield model, we confirm this intuition by simulating “differentiation” through a reduction of the system’s temperature. Subsequently, we develop a potency score based on the intrinsic dimension of expression profiles and confirm its efficacy in capturing differentiation processes, and in distinguishing stem or pluripotent cells from committed cells across various species and biological contexts.

## Results

### The intrinsic dimension of single-cell RNA sequencing data and how to use it

The output of a scRNA-seq experiment is a count matrix reporting the number of detected transcripts from the *D* possible genes in the *N* single cells that have been sequenced. This count matrix naturally defines a set of *N* points in a *D*-dimensional expression space. However, as discussed above, we do not expect that these points can randomly occupy the whole expression space due to regulatory mechanisms and constraints. Therefore, the system should display an intrinsic dimension (ID) lower than the embedding dimension *D*. The ID represents the number of coordinates actually needed to approximately specify the positions of data points with minimal information loss [21]. From a geometric perspective, the data points of several complex systems belong to a relatively low dimensional manifold embedded in the high dimensional data space [15]. In these cases, the ID precisely represent the dimension of the data manifold. The problem of estimating the ID of a dataset has been faced in multiple fields, from physics to computer science, and several ID estimators have been proposed [22–26]. We considered different estimators and analyzed their strengths and weaknesses in the context of expression data. This section summarizes how a robust ID-score can be defined and evaluated on different scRNA-seq datasets.

We selected algorithms belonging to the two main categories: projective and geometric estimators. Projective methods are essentially based on the analysis of the eigenvalues of the *D* × *D* data covariance matrix and aim to extract the minimal number of directions that captures the data variance [25, 26]. Principal Component Analysis (PCA) is an example of a popular linear projective method. On the other hand, geometric or fractal methods leverage on the observation that the density of neighboring points at a fixed distance depends on the ID of the data manifold [22–24].

All ID estimators depend on the sample size, defined by the number of sequenced cells, at least when the system is undersampled (Supplementary Information file, section 1). Given the high dimensionality of the expression space compared to the typical number of sequenced cells (often one or two orders of magnitude less), gene expression datasets are reasonably in the under-sampled regime. In fact, estimators do not converge on the scRNA-seq datasets we analyzed (Supplementary Fig. 1). The convergence behavior of different estimators with the sample size remains poorly characterized, therefore it is not trivial to extrapolate the correct ID.

To address this issue, we compare the estimated IDs of different datasets using the same sample size through random sub-sampling. This method also provides a measure of uncertainty, shown as error bars in the figures, consisting in the standard deviation of ID across sub-samples. As expected for a complex and structured system, the absolute ID values we measure are significantly smaller than the gene repertoire, ranging from units to hundreds depending on the cell populations we consider (Supplementary Tab. 1). These values still depend on the sub-sampling size. However, we are mainly interested in relative values and trends, such as the relative potency of cell sub-populations, the cell potency as a function of time or differentiation stage. Therefore, we can define an ID-score by rescaling the ID measurements in the [0, 1] range (Supplementary Eq. (1)).

As mentioned before, we tested the robustness of the results across ID estimators, but in the main body of this paper we adopt an ID-score based on a specific geometric estimator, called TWO-NN [23, 27]. This estimator is based on the statistics of the distances between each point and its first two neighbours. The focus on such a local property of the dataset makes it more robust to curvature effects and less sensible to outliers with respect to other methods such as PCA. The detailed comparative analysis of different estimators is reported in Supplementary Information file, section 2.

Before analyzing actual biological datasets, we used a toy model to test the basic hypothesis that a system defined by a rugged landscape displays a decrease in ID as the trajectories are progressively constrained by the landscape geometry. We confirmed our hypothesis using the Hopfield model [6] as a testing ground (the detailed analysis is reported in Supplementary Information file, section 3). Although this model was originally developed for memory storage in the brain [28], it has often been used as a possible abstract mathematical formalization of the Waddington landscape [29–32]. We mimicked the differentiation process by progressively freezing the system. The trend of the ID of configurations with temperature confirms our initial intuition (Supplementary Fig. 2).

### The intrinsic dimension decreases with developmental time

As a first test of a link between differentiation and ID, we analyzed a large number of datasets related to embryonic and fetal development across various model organisms. The global level of specialization in the cell population generally increases with developmental time, and this should induce a reduction of the ID. Indeed, Fig. 1 shows a robust and consistent decreasing trend during embryo development of different species (C. elegans, mouse, and zebrafish). The trend spans several developmental time windows and looks progressive, rather than characterized by an abrupt transition at a specific point. In fact, we can observe a decrease in the intrinsic dimension during the first days of mouse gastrulation (Fig. 1B), but also comparing fetal versus neonatal or adult cells (Fig. 1E). Datasets from independent studies on the same biological system, based on different experimental and sequencing protocols (as in the case of Figs. 1A,D), lead to analogous conclusions, thus supporting the robustness of the trend.

**Fig. 1:**
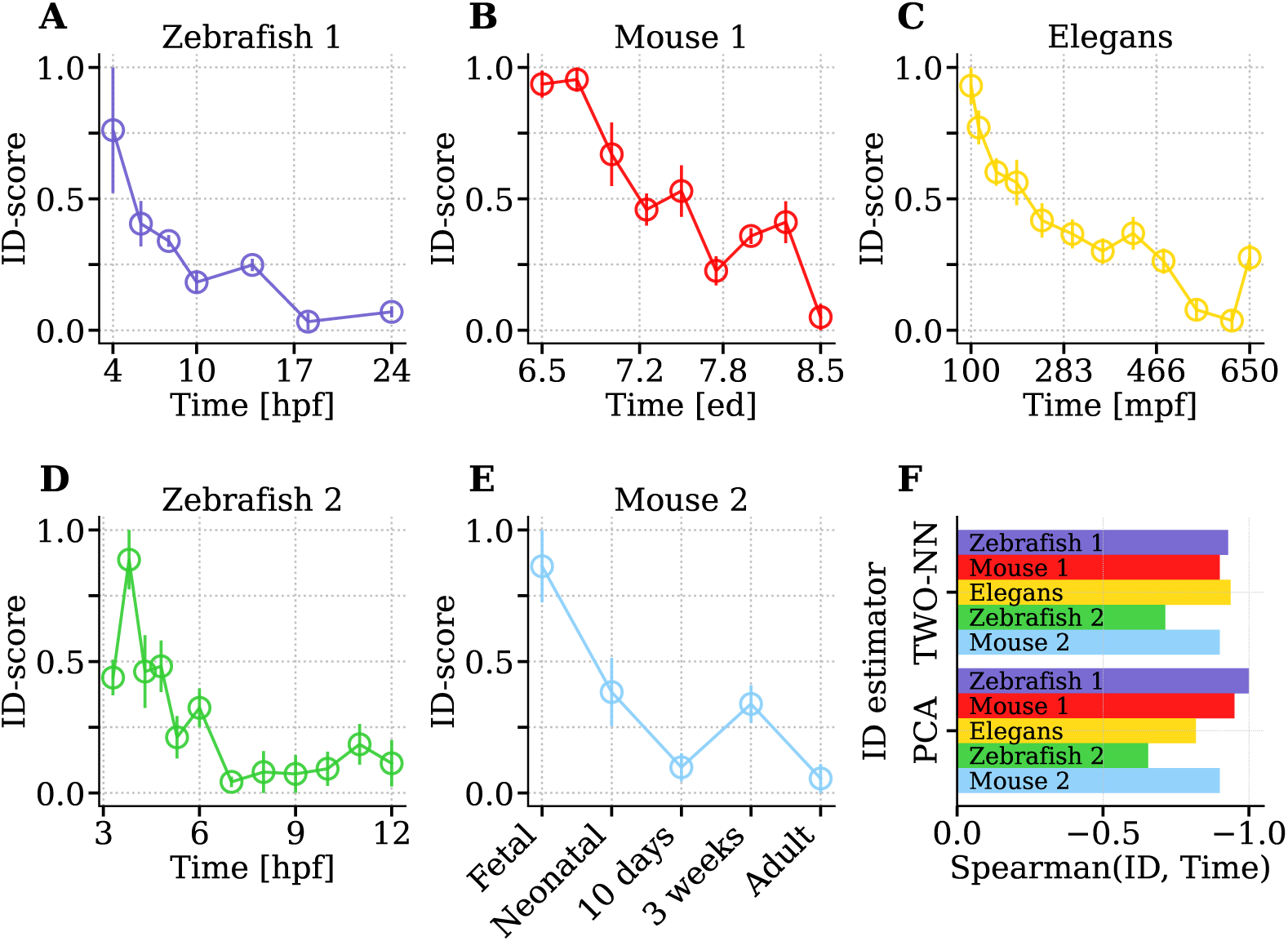
The contraction of the intrinsic dimension of expression profiles during embryo development. ID measurements performed over 10 sub-samplings and re-scaled in the [0; 1] interval (see the Methods section) are reported (circle ± error bar = mean ± standard deviation). Supplementary Tab. 1 specifies the number of cells and genes considered for each dataset. The different panels refer to: **A**) Zebrafish embryogenesis [33], spanning from 4 to 24 hours post-fertilization (hpf); **B**) Mouse gastrulation and early organogenesis [34] between embryonic day (ed) 6.5 and 8.5; **C**) C. Elegans embryogenesis [35] from 100 to 650 minutes postfertilization (mpf); **D**) Zebrafish embryogenesis [36], in the time window 3.3-12 hpf; **E**) Mouse Cell Atlas [37, 38], fetal to adult progression. **F**) Spearman correlation between developmental time and the ID calculated with TWO-NN and a PCA-based estimator, specified in Supplementary Eq. 5.

As motivated in the previous section, the ID-score reported in the figures is based on the TWO-NN estimator. However, the ID dependence on developmental time does not depend on this choice. As an illustrative example, we can consider two estimators based on very different assumptions and on different scales. Specifically, we can compare the local geometric estimator TWO-NN [23] with PCA-based observables (such as the methods described by Supplementary Eqs. (5)(6)(7) in section 2 of Supplementary Information file) that focus on the diversity of the eigenvalues of the whole dataset covariance matrix. Fig. 1F shows that one of these alternative ID quantification indeed leads to similar temporal trends across all different datasets, as confirmed by the high values of the Spearman coefficient. The more comprehensive analysis reported in section 2 of the Supplementary Information file, proves that the agreement between estimators is more general. However, methods based on “local” statistical properties, such as TWO-NN, are less affected by the presence of small sub-populations of cells with radically different profiles, which can strongly affect the measured ID when using projective methods. An example, which we discuss in Supplementary Figs. 3C,D, is the presence of mature red blood cells in the sample. Indeed, these enucleated cells present a highly specific expression profile [39].

The decreasing trend in the ID-score is not a sole property of the whole embryo. It can be equally observed by examining the development of a single organ or tissue, as depicted in Fig. 2. This observation suggests a form of scale-invariance with respect to anatomical resolution. In particular, Fig. 2A and B focus respectively on pancreatic endocrinogenesis and on the formation of the cerebral cortex in mouse. Fig. 2C refers to zebrafish neurogenesis. Fig. 2D provides instead a comparison between fetal and adult stages of seven organs. Despite the significant variability between organs, a consistent decrease in ID after development can be observed. The same behavior emerges when considering artificial embryoids [40] (Fig. 2E). The expression profiles of mouse embryoids at day 8, which completed gastrulation to neurulation and organogenesis, have a clearly lower ID with respect to the less differentiated embryoids at day 6.

**Fig. 2:**
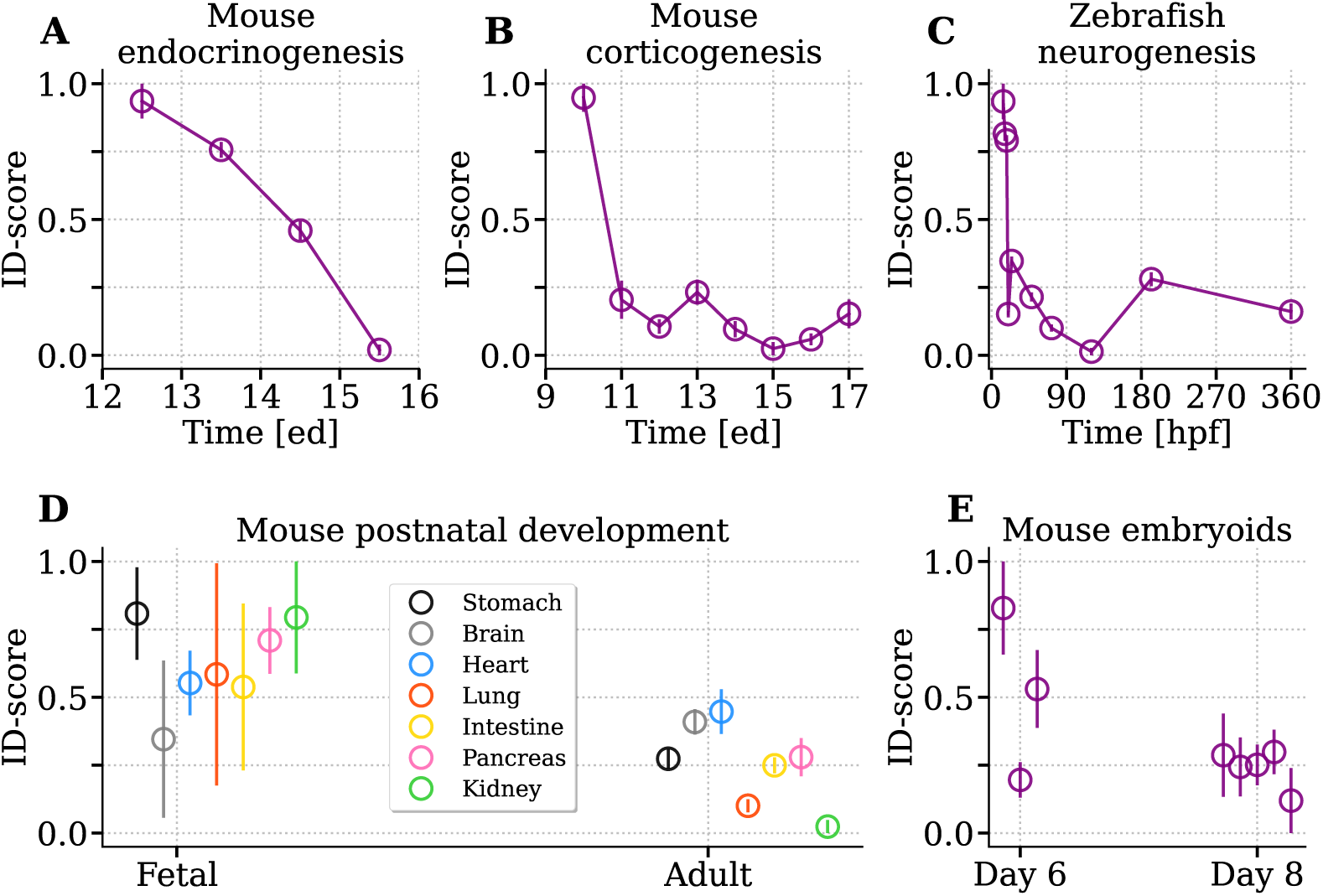
Organogenesis is accompanied by a decrease of expression intrinsic dimension. We analyzed the following organ developmental processes: **A**) Mouse pancreatic endocrinogenes between 12.5 and 15.5 ed [41]; **B**) mouse corticogenesis [42] in the time window 10.5-17.5 ed; **C**) Zebrafish neurogenesis [43] between 14 and 360 hpf. **D**) The Mouse Cell Atlas dataset [37, 38] allows the comparison between the fetal and adult stage of 7 organs. **E**) Finally, embryoid development [40] is analyzed. Three biological samples of embryods cultured for 6 days are compared to five embryoids cultured for 8 days.

In a Waddington landscape scenario, the reduction of the ID should be generally accompanied by an increase in the correlation between genes expression levels, as a result of the geometrical constraints due to gene regulation. As a large-scale measure of the level of correlation, we can consider the size of the gene-gene network whose links represent linear correlations above a certain threshold. The network size grows with differentiation (Supplementary Fig. 4), indicating a global increase of correlation. An analogous trend can be observed by changing the temperature in the Hopfield model.

### The intrinsic dimension reflects the differentiation potential

The observed decrease of the ID with development time can be ascribed to two main possible factors: the overall progressive differentiation of the cell population and the proliferation in the number of the cell types during tissue and organ formation. There is indeed a possible relation linking the measured ID and the number of cell types, which typically grows during embryogenesis (Supplementary Fig. 5).

In a Waddington landscape scenario, cell types are represented by different basins of attraction that can have specific geometries and intrinsic dimensionality, depending on the level of complexity and the degree of gene regulation defining each cell type. Therefore, the expression profiles of cells composing an organ or a whole embryo are expected to lie on a structured and composite manifold. ID estimators are affected by this heterogeneity. Specifically, as we show in detail in section 2 of the Supplementary Information, when the data points belong to a composition of manifolds with heterogeneous dimensions, the ID estimate given by TWO-NN is dominated by the low-dimensional manifolds. Therefore, the simple increase in the number of cell types, which typically occurs during development, can induce a trend in ID. In fact, there is an increasing chance of observing a cell type associated to a low dimensional manifold if their number increases and they have heterogeneous IDs (Supplementary Figs. 6 7).

Since we are interested in evaluating the ID as a score for cell potency, we need to disentangle this spurious effect and quantify the actual correlation between ID and differentiation level. To this aim, we collected several well established and annotated differentiation trajectories. Along these trajectories, cell types can be roughly ordered by their differentiation level, and we can test if the ID can recapitulate this ordering without relying on prior biological information.

As a first example, the process of pancreatic endocrinogenesis in mouse is known in sufficient detail to draw the diagram of the lineage relationships between pancreatic cells that summarizes the differentiation process [41]. Fig. 3A shows that the ID-score can correctly order the cell types in terms of their potency along d differentiation lineages, correctly reproducing the known relationships between cell types only from data geometry.

**Fig. 3:**
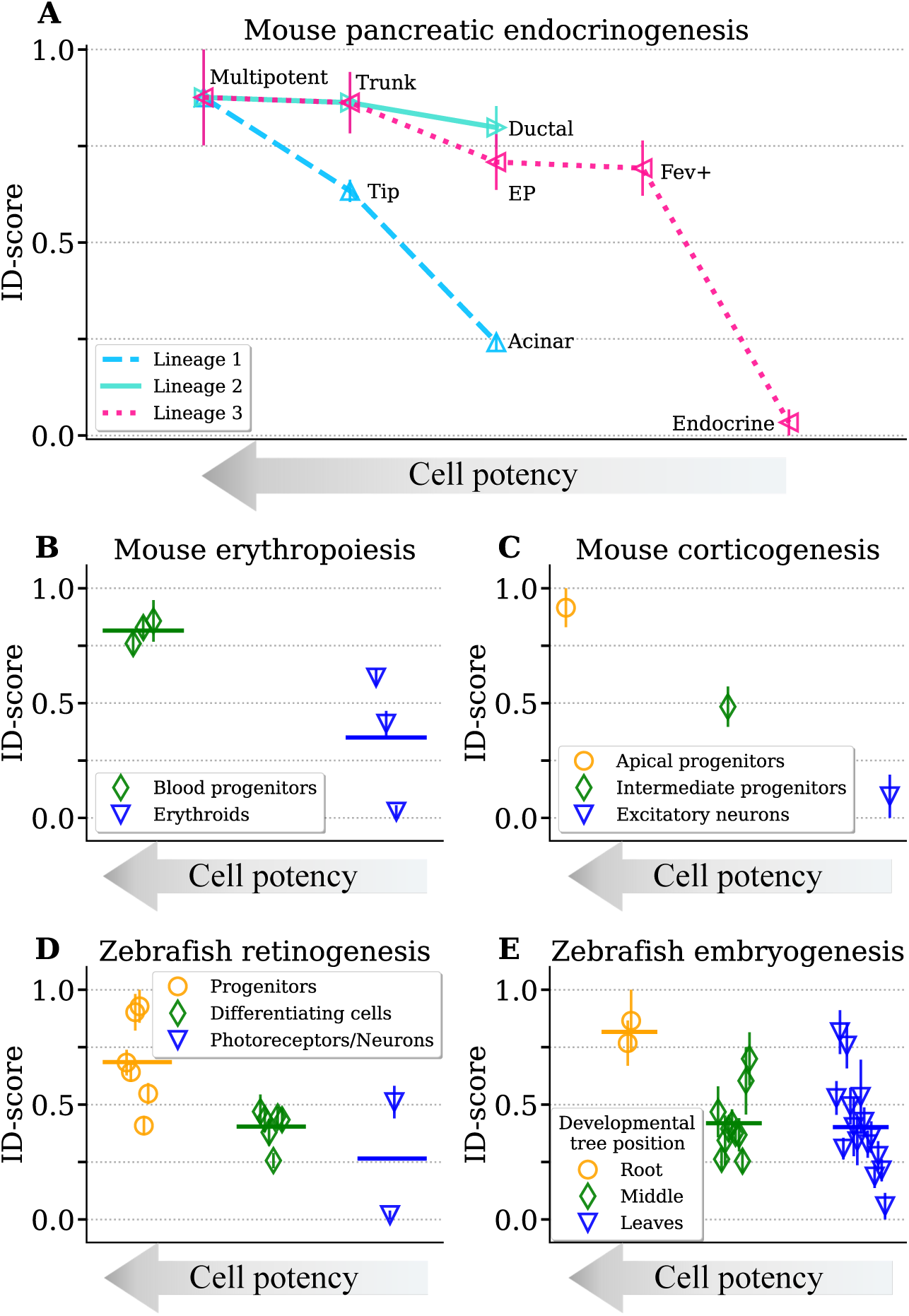
The intrinsic dimension reflects the potency of cell types along developmental lineages. **A**) The hierarchy of cell types along three main differentiation lineages of pancreatic endocrinogenesis (as reconstructed in ref. [41]) can be well reproduced by the intrinsic dimension. **B**) In the production of mouse erythroids [34], the ID-score can distinguish the class of haematoendothelial and blood progenitors from erythroids. Horizontal lines show the average ID-score of cell types belonging to a similar potency level. **C**) In mouse corticogenesis [42], we can correctly order apical progenitors, intermediate progenitors and excitatory neurons. **D**) In the formation of retinal neurons in zebrafish [44], different cell types can be group in the broad classes of retinal progenitors, differentiating retinal cells and retinal neurons/photoreceptors. The ID-score can well separate their different levels of differentiation. The specific cell types are reported in Supplementary Information, section 6. **E**) Cell types from zebrafish embryos are roughly ordered according to their position in the developmental tree reconstructed in ref. [36].

The developmental lineage relative to erythroid cell formation during mouse gastrulation has also been reconstructed in detail [34]. In particular, cell types can be ordered according to their potency from progenitor cells to the final erythroids. Fig. 3B demonstrate that the ID can clearly separate the cell types belonging to these two classes. Similarly, the apical and intermediate progenitors can be compared to excitatory neurons, a fully differentiated cell type, in the process of mouse corticogenesis [42]. Also in this case, the ID score shows that the progressive differentiation corresponds to a reduction of the dimensionality (Fig. 3C).

An analogous analysis can be performed on different species to test the robustness of the results. Zebrafish embryogenesis is a well studied system in which the cell types have been characterized and can be ordered by potency along differentiation lineages. In particular, looking at the expression of well-established gene markers, Farnsworth et al. annotated three main cell clusters with increasing level of specialization: retinal progenitors, differentiating retinal cells and final retinal neurons and photoreceptors [44]. Fig. 3D shows how the ID correctly reproduces this potency ranking.

Finally, in reference [36], an alternative diffusion-based computational framework was used to infer the developmental trajectories in zebrafish embryogenesis. Each cell type can thus be placed on a tree-like structure. The tree root, corresponding to pluripotent cells, and the different branch annotations were validated by marker gene expression patterns. Starting from these inferred lineages, we can distinguish three groups of cell types of decreasing potency level: the cell types close to the tree root, the intermediate cell types along the branches, and finally the differentiated cell types at the tree leaves. Fig. 3E reports the coherent decrease of the ID along the differentiation tree.

The ID seems to recapitulate the potency trends in dynamic differentiation processes related to embryo development and organogenesis. However, a general measure of potency should be able to discriminate between pluripotent and differentiated cells also during tissue homeostasis, maintenance and regeneration. The small freshwater Hydra polyp is specifically studied because of its continual self-renewal and regeneration ability [45]. In this system, stem cells have been separated from fully differentiated cells belonging to endodermal epithelial, ectodermal epithelial, and interstitial lineage, using a combination of computational tools and prior knowledge based on marker genes and spatial localization [45]. The ID of genome-wide expression profiles can clearly capture the potency difference between stem and specialized cell types in all three lineages (Figs. 4A,B,C).

**Fig. 4:**
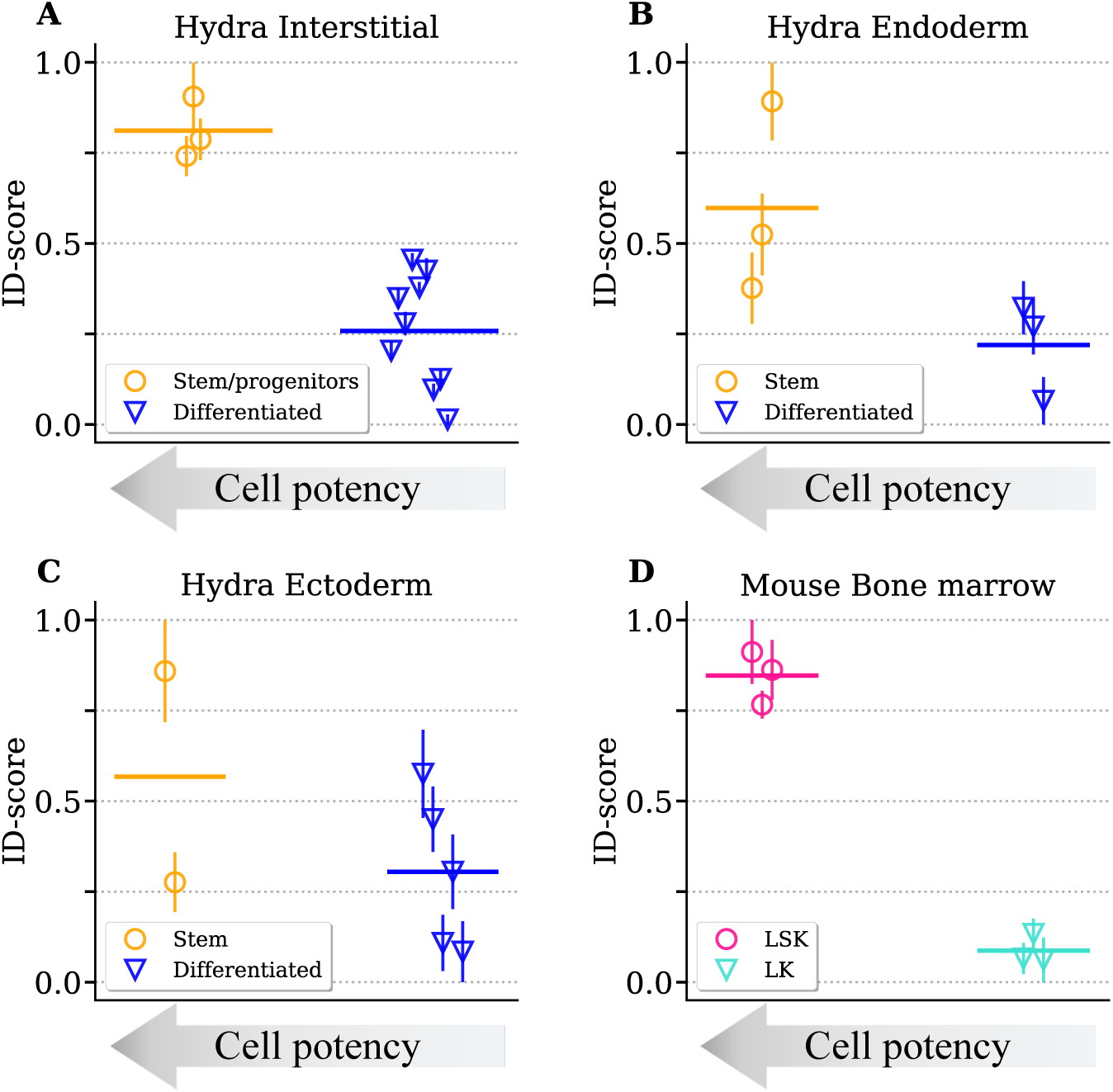
The intrinsic dimension can discriminate between stem cells and differentiated cells in tissue turnover. The ID-scores relative to clusters of stem/progenitor cells versus differentiated cells are compared for the three hydra epithelial layers: interstitial (**A**) endoderm (**B**) and ectoderm (**C**)(data from [45]). The specific cell types corresponding to each point are reported in Supplementary Information, section 6. **D**) The ID-score of two cellular populations (LSK, LK) isolated in hematopoietic stem cells [46] is reported. Orange points refer to the immature LSK cells, while blue points to the more differentiated LK cells.

As a final example, we consider hematopoiesis, i.e., the process of continual turnover of blood cells driven by haematopoietic stem cells in the bone marrow. A large-scale study in mouse identified two sub-populations (LSK and LK) with a different level of maturity in hematopoietic stem and progenitor cells using a sorting procedure based on two markers [46]. LSK cells are early progenitors, not committed to a specific blood cell lineage, and express high values of Sca-1 (stem cell antigen-1) and c-Kit (a transmembrane receptor associated to immature state). On the other hand, LK cells exhibit low levels of c-Kit, indicating an intermediate state of maturity. Once again, the ID can well discriminate the different differentiation potentials of the two sub-populations (Fig. 4D).

## Discussion

The process of cell differentiation is a comprehensive reorganization of cellular physiology. The robust and consistent differentiation patterns observed at the macroscopic level are ultimately determined by a complex orchestration of a high number of molecular processes heavily subjected to stochasticity at the single-cell level. In analogy to statistical physics systems, an ordered macroscopic state emerges from a high-dimensional and noisy microscopic behavior [47]. Therefore, even if key specific regulators play a dominant role in differentiation [48–50], large-scale observables should more reliably capture the extensive molecular reprogramming involved in differentiation with respect to the inherently noisy expression of a handful of “marker genes” [51]. Modern scRNA-seq techniques offer an access to such genome-wide observables.

Taking inspiration from the Waddington landscape picture, we have shown that indeed global geometrical properties of single-cell expression profiles are indicative of the differentiation level, which can be robustly recapitulated by a single score based on the intrinsic dimension. Importantly, this ID-score does not require specific or complex data pre-processing or prior biological knowledge about the system. For instance, the reported ID trends are robust to common gene selection techniques (Supplementary Fig. 8) and consistently emerge across species and tissues. This simplicity and robustness of the ID-score allows its straightforward integration in current analysis pipelines [10]. If different sub-populations are identified in a sample, for example through clustering or thanks to known relevant molecular players, the ID-score can order these cell groups along a potency line.

A plethora of computational methods, often called pseudotime inference tools, try to align single cells along trajectories using the similarity of their expression profiles [36, 52–55]. These trajectories should reflect continuous biological processes such as developmental paths, although inferred from static expression snapshots. However, the vast majority of these methods need prior information about the identity of the starting (or end) cells, thus ultimately about the differentiation direction [52]. Our ID-score can robustly provide the correct ordering of cell groups or equivalently the pseudotime direction, thus constraining the likely differentiation diagrams.

More generally, time clearly plays a crucial role in development. However, the order and tempos of differentiation steps are not conserved across species and systems [56]. Therefore, the relationship between time and differentiation potential could be arbitrarily complex and system-dependent. As RNA sequencing becomes more and more precise, our quantitative measure of potency can be used to estimate the actual rate of differentiation over time, unlocking the possibility of quantitative comparisons, for example of the speed of embryo or tissue development across different species [57] or between natural and laboratory models such as organoids [40, 58]. The analogy between the Waddington landscape and statistical physics systems has long been recognized. Consequently, tools and ideas from statistical mechanics have percolated into single-cell analysis [14, 59]. Specifically, measures based on the entropy of expression profiles have been proposed as proxies for stemness [60–65]. The basic idea stems from the observation that high variability in expression profiles is often associated with stem cells, and entropy is a theoretically grounded measure of variability that goes beyond simple variance-based evaluations [60]. While entropy seems to capture known differentiation trends in some datasets, it fails in several instances where the ID-score robustly recovers known potency hierarchies (Supplementary Fig. 9C,F). These two quantities capture different, and in principle, independent aspects of the dataset statistics, and thus can be jointly used to extract global patterns in expression profiles. However, entropy values depend on the number of available states of the system, which is set by the number of detected genes. Unfortunately, the sparsity of scRNA-seq datasets is strongly influenced by technical noise due to the sampling process of RNA sequencing [17]. This effect suggests a higher robustness of geometry-based measures like intrinsic dimension. In fact, to enhance robustness, many proposed entropy-based tools integrate additional information, which may not always be available, such as the protein-protein interaction network [65] or gene functional annotations [63].

Similar considerations hold true for another proposed tool for potency estimation, which leverages directly on the total number of expressed genes [66]. Transcriptional diversity often correlates with potency, probably because differentiated cells selectively switch-off certain pathways. However, this measure does not always capture potency trends in the datasets we explored (Supplementary Fig. 9B,E), possibly due to its high sensitivity to sampling noise.

Defining a measure that can quantitatively and robustly capture the potency level of a cell population is useful beyond the reconstruction of natural developmental trajectories. For example, cell reprogramming experiments promise to have relevant applications from regenerative medicine to disease modelling and drug testing [49, 67, 68]. In reprogramming protocols, the goal is to induce pluripotency in differentiated cells. A quantitative potency measure, such as the ID-score, can complement existing methods by providing a marker-free metric to identify cell subpopulations that have successfully achieved pluripotency and to assess the extent of their potency.

Finally, the Waddington landscape metaphor has also been invoked for understanding cancer etiology and progression [69]. This parallel suggests that our approach could represent a useful quantitative tool in this context as well.

## Methods

### Datasets

We collected recently published scRNA-seq datasets that are readily accessible through the GEO repository [70] or other online repositories. These datasets span various model organisms, and describe embryonic development, organ/tissue development or specific differentiation lineages. The experiments were performed in independent laboratories with different protocols. However, we only selected experiments using Unique Molecular Identifiers (UMIs). The full list of datasets is reported in Supplementary Tab. 1 and briefly described in the Supplementary Information, section 6.

### Data pre-processing

Cells were filtered according to three criteria based on thresholds on the total number of reads, the number of detected genes and the mitochondrial percent. We directly applied to each dataset the thresholds reported by the authors (Supplementary Information, section 1). When applicable, doublets identified by the authors were also excluded.

We only considered protein-coding genes as annotated by the data mining tool BioMart, accessible via the Ensembl database [71]. A complete list of proteincoding genes is not available for Hydra. Therefore, in this case, we used all the genes contained in the dataset.

A relevant step in most analysis pipelines of scRNAseq data is the normalization introduced to partially compensates for heterogeneity in sequencing depth [10]. We simply divided the gene transcript counts in a cell by the total number of detected transcripts, defining cell transcriptional profiles with relative abundances.

### Intrinsic dimension estimators and the ID-score

We considered different ID estimators. Among the class of projective or PCA-based methods, we used three different statistical properties of the covariance matrix, all related to the data ID: the number of principal components *d_PCA_* required to explain a given percentage of the total data variance (Supplementary Eq. 5); the entropy of the normalized covariance eigenvalues *H_PCA_* (Supplementary Eq. 6) and the complementary of their Gini coefficient *G_PCA_* (Supplementary Eq. 7).

In the class of geometric/fractal methods, we selected the TWO-NN estimator [23] and the FCI estimator [24], respectively relying on a local and a multiscale approach. The results reported in the figures are based on TWO-NN. However, the reported trends are generally very robust to the estimator choice (Supplementary Figs. 10 11 12 13). A more detailed comparative analysis and a mathematical description of the estimators are reported in the Supplementary Information, section 2. All ID estimators have a specific dependency on the sample size and on the dimension of the embedding space, i.e., the number of cells and genes in the count matrix (Supplementary Fig. 1). To remove the dependency on the sample size, we randomly sub-sampled a number of cells corresponding to 75% of the least represented cell group (with an upper bound of 5000 cells for computational reasons) and measured the intrinsic dimension with different estimators over 10 independent sub-samplings. The mean and the standard deviation of these values are reported as circles and error bars in the figures. Analogously, we considered the same number of protein-coding genes for all cell sub-populations that have to be compared. The exact number of cells and genes used are dataset-specific and are reported in Supplementary Tab. 1. Since the absolute ID value depends on non-biological variables such as the sample size, we decided to define a scaled ID-score, whose values are in the [0 − 1] range, to be compared with the potency level (Supplementary Eq. (1)).

## Declarations

### Ethics approval and consent to participate

This study utilized publicly available single-cell RNA-seq data from E-MTAB repository and Gene Expression Omnibus database. As such, no ethical approval or informed consent was required.

### Consent for publication

Not applicable.

### Availability of data and materials

The scRNA-seq datasets we analyzed (Supplementary Information file, section 6) are publicly accessible through the links provided in the corresponding papers. The original code is available at: https://github.com/BioPhys-Turin/The-int rinsic-dimension-of-gene-expression-during-cell-differentiation. Additional data requests can be directed to the corresponding author (M.O).

### Author contributions

M.O. conceived and supervised the study; M.O., M.B., and N.C. identified key results and developed the methodologies; M.B., N.C., and S.L. collected and processed the datasets; M.B. and N.C. analyzed the data; M.C and F.V. worked on the analogies with the Hopfield model; M.O., N.C, and M.B wrote the paper. All authors approved the final draft of the manuscript.

## Acknowledgements

We are grateful to M. Cosentino Lagomarsino and A. Scialdone for useful feedback on our work.

## Appendix

### 1 Data pre-processing and intrinsic dimension estimation

#### 1.1 Filtering and pre-processing

In order to reduce the technical noise due to the sampling procedure involved in RNA sequencing, single-cell expression data are usually normalized [17, 72]. We simply normalized the transcript count 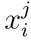 of gene *j* in a cell *i* by the total number 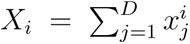 by a vector of relative transcript abundances in the expression space. While this procedure has the goal of eliminating the effect of sampling depth on expression variability, it can also remove relevant biological signals. In fact, the heterogeneity of *X_i_*values is also related to physiological features, such as cell size. With the technical development of sequencing protocols, if biases in the number of detected transcripts become negligible, it would be possible to remove the normalization step and account for the effect of the total number of transcripts on data geometry.

We adopted the same cell quality control suggested by the authors of each open access dataset we analyzed. These criteria typically include filters based on the number of detected genes, the total number of transcripts, in addition to exclusion of putative doublets.

We focused only on protein-coding genes following the annotation available on BioMart, a data mining tool accessible via the Ensembl database [71]. The only organism for which a list of protein-coding genes is not available is Hydra. In this case, we used all the transcripts contained in the dataset.

We intentionally minimized the pre-processing procedure to test the robustness of our method, to make it widely applicable and to avoid the introduction of arbitrary data transformations such as complex normalizations, data imputation techniques or non-linear data projections [10]. The hypothesis is that relevant global geometrical properties and trends of the data can be extracted from essentially raw data and could instead be confounded by complex pre-processing procedures.

However, our analysis should not be dramatically affected by standard pipelines, and it could even benefit from data corrections able to attenuate confounding factors such as batch effects [73]. In fact, Supplementary Fig. 8 shows an example of how the temporal trend of the ID-score reported in Fig. 1E changes when a standard feature selection technique is applied. Specifically, we selected only the highly variable genes following one of the most popular gene selection strategies [74]. The estimation of the ID-score is robust with respect to this choice, and the variability between different sub-samplings (error bars) is reduced.

#### 1.2 Dependence of intrinsic dimension estimations on the sample size and the sub-sampling procedure

As discussed in the main text, the estimated values of intrinsic dimension depend on the sample size in the under-sampled regime. Supplementary Fig. 1A reports this dependence for the TWO-NN estimator applied to two simple synthetic datasets. These datasets are obtained by sampling from a 50-dimensional and a 80-dimensional hypercube, respectively, embedded in a 1000-dimensional space. The true ID value is underestimated across several order of magnitudes of the sample size because of the curse of dimensionality [9].

In the case of real datasets, in particular of scRNA-seq count matrices, even if we do not know the true ID, we can observe the relationship between the estimated ID and the number of data points (Supplementary Fig. 1B). The trend of the curves suggest that we are far from saturation and thus the estimated values still crucially depend on the number of cells available. Fortunately, the primary goal of our analysis is not the exact estimation of the ID. We are mainly interested in the evolution of this quantity during development and differentiation, and in ranking cell populations in terms of potency. To make these ID comparisons meaningful we need to compare the ID of groups with the same number of cells. To this aim, we measure the ID on random subsets of fixed size extracted from each cell cluster. With this procedure we obtained Figures 1 2 3 4, where the average ID values and their standard deviations across independent sub-samplings are reported. To determine the size of each sub-sample, we identify the least represented cell cluster and consider the 75% of its number of cells (see column 3 of Supplementary Tab. 1). For computational reasons, we imposed an upper bound of 5000 cells to the size of the sub-samplings. The number of sub-samplings is set to 10 for every trend we report. Clearly, the variability introduced by the sub-sampling procedure is affected by the initial size of each cluster. The less populated clusters will often exhibit the lowest variability as measured with the standard deviation. The absolute ID values measured with the TWO-NN method, although influenced by the undersampling, are significantly lower (∼ 10^1^) than the number of genes present in the original library (∼ 10^4^).

#### 1.3 Definition of a normalized ID-score

Since the absolute ID values have a technical dependency on the sample size, we decided to define a re-scaled ID-score and test its correlation with the potency level. Specifically, the measured ID values are normalized in the [0 − 1] range, by defining

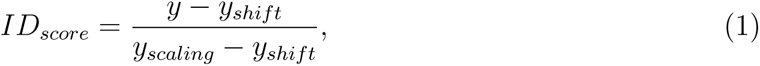

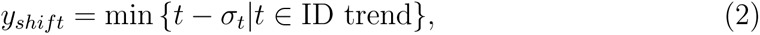

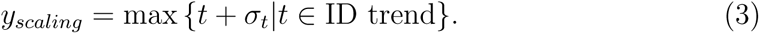

The *y* value corresponds to the average estimated ID (with the chosen estimator) over the random sub-samplings. For every dataset, we subtract the minimum ID value *y_shift_* and we scale the values with the maximum value *y_scaling_*. The values of *y_shift_* and *y_scaling_* are computed taking into account the variability across subsamplings. Therefore, the ID-scores in the paper figures are constrained in the interval [0 − 1] with the error bars included. The values of these parameters for each dataset are reported in Supplementary Tab. 1.

### 2 A comparison of intrinsic dimension estimators for scRNA-seq data

#### 2.1 Tested ID estimators

Among the plethora of intrinsic dimension estimators, we can first distinguish between global estimators and local estimators. Local estimators focus on data neighborhoods of a given scale, while global estimators look at the geometry of the whole dataset. An alternative distinction can be made between projective, and fractal or nearest neighbor-based estimators.

We tested estimators belonging to different classes and relying on different assumptions. Specifically, TWO-NN is local and belongs to the class of nearest-neighbors methods. FCI is a multiscale approach and can be classified as a fractal method. Finally, PCA-based ID estimators are global quantities based on linear projections.

- The TWO-NN algorithm has been proposed in [23] and is based on the computation of the typical distances among neighboring points (cells) in the data space. In particular, the method reconstructs the distribution of the distance ratios *µ*. For each data point, the value *µ* = *r*_2_*/r*_1_ is the ratio between its distance to its second (*r*_2_) and first (*r*_1_) neighbor. Assuming local data density homogeneity and considering the sampling process as a Poisson point process, Elena Facco et al. demonstrated that *µ* follows a Pareto distribution *f* (*µ*):

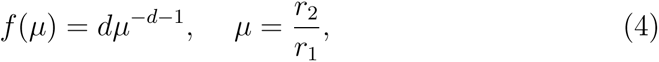

with an exponent defined by the intrinsic dimension *d*. The ID is inferred from a fit on the curve {log *µ,* − log (1 − *F* (*µ*)}, where *F* is the cumulative function of *f*. To make the fit robust to outliers we discard the highest values of *µ* and consider only the remaining 90% of values, as suggested by the authors of [23]. Due to the normalization (see Supplementary section 1.1), our data points have non-integer coordinate values. The metric we used to compute the distances is the euclidean distance. The Manhattan distance could be more appropriate if one decide to use directly the integer transcript counts.
- The Full Correlation Integral (FCI) [75] is a fractal method that has been recently proposed as a development of the classic Grassberger and Procaccia method [22] for undersampled datasets.
- PCA-based observables. Given *N* cells, *D* genes and a counts matrix ***X_N×D_***, the eigenvalues ***λ*** = (*λ*_1_*, λ*_2_*, …, λ_m_*) (where *m* = min {*N, D*}) of the covariance matrix ***C_D×D_*** specify the contribution to the total variance of the data given by each eigenvector or principal component. The relative magnitude of these values contains information about the intrinsic dimension of the dataset. We can count how many components *d_PCA_* are needed to retain a certain percentage *V*_th_ of the total data variance *V*_0_, and use this value as a proxy for the ID. In other words, we can define:

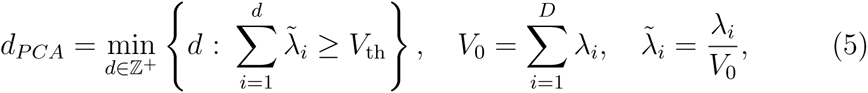 With this definition, the absolute ID value depends on an arbitrary parameter *V*_th_, but again we can define a normalized ID-score. Alternatively, we can introduce two other methods of evaluating the non-uniformity of the set of eigenvalues ***λ***, with the general idea that the level of heterogeneity in the eigenvalues increases as the ID decreases. Specifically, we defined two measures based on the Shannon entropy (*H_PCA_*) and on the Gini index (*G_PCA_*) following the expressions:

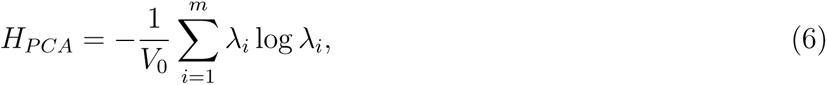

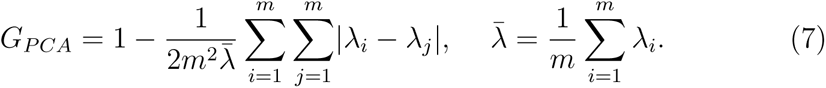 These two quantities mainly differ in the weight attributed to small eigenvalues. The entropy-based index of Supplementary Eq. (6) is generally more sensitive to them. PCA-like methods are linear methods. Therefore, if the true data manifold has a curvature, ID estimations based on linear projections will systematically overestimate the ID. Finally, PCA-based method are reliable only if the number of samples is greater than *d* log(*d*), with *d* being the (usually unkown) intrinsic dimension [24].

#### 2.2 How the presence of multiple manifolds with heterogeneous dimensionality affects ID estimators

Single-cell data are characterized by strong heterogeneity having both biological and technical origin [17]. A large source of biological variability in the datasets we analyzed is the presence of multiple cell types (Supplementary Fig. 5). Since each cell type is characterized by a specific gene expression program and set of regulations, the geometry of expression profiles of cells belonging to different cell types could be radically different and characterized by different intrinsic dimensions. In other words, the points of our dataset could be partitioned on different manifolds with specific geometrical properties and dimensions, embedded in the gene expression space. However, most ID estimators implicitly assume the presence of a single true ID that have to be estimated [76]

In this section, we analyze what is the effect of having a composition of manifolds of different dimensionality on the ID values that can be obtained with TWO-NN and PCA-based estimators. In particular, we consider the illustrative case of points distributed over two manifolds of different dimensions using synthetic (Supplementary Figs. 3A,B) and real data (Supplementary Figs. 3C,D). The main result is that all estimators do not report an average or intermediate value between the true IDs, but are generally biased towards low dimensional values.

- **PCA-based methods.** If a dataset is composed of two clusters that are well-separated in the gene expression space, the direction of separation could be detected as the first principal component, as it captures a large variance. Consequently, the covariance matrix will have a very large first eigenvalue, which can lead to a biased estimation of the ID towards low values (see Supplementary Eqs. 5 6 7), regardless of the actual ID of the two clusters. This is clear in the examples in Supplementary Fig. 3A and C. The first principal component is the direction along which the two clusters of red and blue points separate. More quantitatively, if we consider separately erythroid and primitive streak cells (Supplementary Fig. 3C), we estimate respectively *d_PCA_*=3 and *d_PCA_*=381 (with threshold *V*_th_=0.9). On the other hand, if we consider the dataset as a whole, we estimate *d_PCA_*=2. The pronounced separation of expression profiles of these two cell types is probably due to the fact that erythroid cells do not have nuclei and organelles. The presence of such cell “outliers” can bias the PCA-based estimators towards low ID values.
- **TWO-NN.** In the TWO-NN method, points lying on low-ID manifolds have a greater impact on the overall estimated ID. The method relies on the distribution of the distance ratios *µ* (Supplementary Eq. (4)), where the right tail (i.e., large values) is dominated by points sampled from low-ID manifolds. Intuitively, in high dimensions, neighboring data points are more likely to be at similar distances. An example of this effect is reported in Supplementary Fig. 3B and D. The two plots show the cumulative distribution *F* (*µ*), which is used to estimate the ID by TWO-NN. This distribution is evaluated either on the whole dataset or separately on the two manifolds (two hypercubes in B or two different cell types in D). The global distribution is much closer to the one corresponding to the low-dimensional part of the dataset. Therefore, in the presence of manifolds of different dimensions, TWO-NN estimates on the whole dataset are strongly biased towards low dimensions. On the other hand, thanks to its local nature, TWO-NN exhibits better tolerance for isolated outliers compared to PCA-based estimators.

These considerations can be extended to the case of many cell types. The ID can naturally drop if the number of cell types in the dataset increases and the cell types are associated to heterogeneous IDs. In fact, if we build artificial datasets by randomly assemble data relative to different cell types, we can observe an average decrease of the ID with the number of cell types (Supplementary Figs. 6A,B)

In several datasets related to development, the number of cell types naturally increases with time following differentiation lineages. Therefore, the ID decrease we observe in time can be due to cell differentiation, but also to the growth in the number of cell types. In fact, the ID-score is generally correlated with this quantity in developmental datasets (Supplementary Fig. 6C). Precisely to separate the two contributions and establish the ID-score as a measure of potency, we also analyzed single differentiation lineages following a single cell type as it differentiates.

#### 2.3 ID across scales: from local to global

As discussed in the previous section, the heterogeneity of the data manifolds can bias the different estimators in different ways. For example, single outliers can bias global PCA-based estimators towards low dimensional values. On the other hand, fractal and local methods such as TWO-NN are more robust to outliers but still sensitive to dimensional heterogeneity.

To better understand the role of the scale (number of neighbor points considered) on the ID estimation in presence of heterogeneous datasets, we introduce a local PCA estimator *Ld_PCA,l_*. The parameter *l* sets the observation scale through the number of neighbor points used to evaluate the covariance matrix. With this local covariance matrix, we can apply the *d_PCA_* method (Supplementary Eq. 5) to estimate the ID. By changing the scale *l*, we can highlight and better understand some critical aspects of the ID estimators in presence of heterogeneous datasets. In general, considering too few data neighbors can lead to strong undersampling and noisy covariance estimates, while using too many neighbours may cause a drop of the ID, due to the influence of outliers or to the presence of multiple cell types (Supplementary Fig. 3 and 6).

Supplementary Fig. 7 shows in detail how the scale affects the ID estimations on one illustrative dataset from mouse gastrulation [34]. We randomly selected 50 cells in the dataset and their first *l* neighbours (fixing the scale). We then computed the intrinsic dimension of these 50 cell groups and examined their distribution. At very small scales, the *d_PCA_* increases with *l*, just because we are in an extremely undersampled regime and the ID increases with the number of points considered, as in Supplementary Fig. 1. At intermediate scales we start to observe the separation of cell types, and thus a broad and multimodal distribution of IDs. At these scales, the neighborhood of a cell is typically composed by cells of the same cell type. Here, around *l* = 411, we can appreciate that cells of the caudal epiblast and mesoderm show a high intrinsic dimension, while the erythroids have a smaller ID with respect to haematoendothelial and blood progenitors. This ranking largely reflects the known level of specialization of those cell types.

At large enough scales, the data neighbors contains cells of different cell types and the value of local *Ld_PCA,l_* progressively collapse on the global estimator *d_PCA,l_* as expected.

Since in the dataset are present different cell types, the globally estimated ID converges to a value biased towards the low dimensional values, as explained in Supplementary Section 2.2.

#### 2.4 Comparison between ID estimators

This section reports the results of the tests of robustness of our main results with respect to the choice of the ID estimator. In particular, we evaluated the correlation between the ID-score defined with several estimators (Supplementary section 2.1) on each dataset analyzed for the Figures 1, 2, 3 and 4. We also introduce another local estimator *LG_PCA,l_*, which corresponds to a local version of *G_PCA_* (Supplementary Eq. (7)), with a parameter *l* setting the observation scale, i.e., the number of neighbor data points used to evaluate the covariance matrix. We explored several values of *l* to understand effects related to the observation scale.

Supplementary Figures 10, 11, 12 and 13 show how the ID-based rankings obtained with different estimators are generally remarkably conserved. Some disagreement is present in only few cases relative to temporal trends, that are characterized by a strong heterogeneity in cell type composition (Supplementary Fig. 5). As discussed above, some estimators are quite sensitive to the number and heterogeneity of data manifolds, thus explaining the reduced robustness of results in these cases. Supplementary Fig. 11B constitutes an illustrative example of this effect. In this case, we verified by projecting the data with PCA, the presence of a small number of outliers well separated from the bulk of the dataset in the gene expression space. Therefore, global estimators based on PCA are expected to strongly underestimate the ID. As a proof, we can observe the agreement between local estimators (TWONN and *LG_PCA,_*_120_), while more global estimators (FCI or PCA-based observables) lead to different rankings. However, when we get rid of this heterogeneity by manually removing those outliers the agreement between estimators can be restored. In fact, when we apply our analysis to single cell types, the estimations based on different methods are robustly correlated (Supplementary Fig. 12 and 13).

### 3 The Hopfield model as a toy model of the Waddington landscape

As discussed in the main text, we selected the Hopfield model [28] to preliminary test our hypothesis because it shares some key features with the Waddington landscape metaphor. Specifically, the Hopfield model is characterized by an energy landscape with attractors, which correspond to stored memories in the original model. These state attractors can be associated to the typical expression profiles of various fully differentiated cell types in our analogy. In contrast, a stem or multipotent cell is not constrained within a single attractor. Instead, the differentiation process progressively aligns its expression profile with that of a committed cell type. Accordingly, temperature in the Hopfield model can be used as a parameter to regulate the level of “differentiation”. By decreasing the temperature, we can mimic the differentiation process, akin to increasing the number of geometric constraints on the configurations. The key question is whether the intrinsic dimension can quantitatively capture this “differentiation process” in the Hopfield model. In other words, if the intrinsic dimension of an ensemble of spin configurations has a specific increasing trend with temperature.

The correspondence between the Hopfield model and the Waddington landscape has been often invoked, and used to build computational tools for transcriptomic data [29–32, 62, 77, 78]. However, we do not claim that the Hopfield model provides a direct, accurate, and quantitative description of the epigenetic process of cell differentiation. Instead, we aim to test the hypothesis that the ID can well capture the progressive increase of geometrical constraints as the temperature decreases in a model with a complex landscape where all the relevant parameters are under control. To make the analogy more precise, a spin configuration corresponds to a singlecell transcriptomic profile, although the expression levels are discretized to −1 or 1 in the classic Hopfield formulation. The number of spins in our model (*D*) is chosen to be comparable to the number of genes typically present in RNA-seq dataset analysis (*D* ≃ 1 000). The number of trajectories we simulate (*N*) corresponds to the number of cells in a scRNA-seq experiment. For example, *N* = 1 500 in the experiments reported in Supplementary Fig. 2. We tried to reproduce the highdimensional nature of our datasets in the analogous Hopfield model to more reliably check our expectations about the intrinsic dimension, since intuition often fails in high-dimensional settings.

The *P* attractors *ξ^µ^* (the memories of the Hopfield model) are set as random spin configurations. These memories represent the “archetypal” transcriptional profiles of fully differentiated cell types. In the toy model these patterns are random configurations with no particular structure or mutual relations, as it is instead probably the case for transcriptional profiles.

The weights defining the couplings between spins are set by the rule: *W_ij_* = 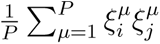. The energy of a state *S* is defined by *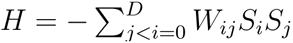*.

The Hopfield model has a critical value *α_c_* for the capacity *α* = *P/D* [7]. When the number of stored patterns is below the critical capacity *α_c_* = 0.138 (and the temperature is below 1), the stored patterns are stable attractors in the energy landscape. We are interested in this regime as an analogous of the Waddington landscape. Given our parameter setting, we have *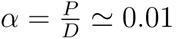*, so we are indeed in the correct region *α < α_c_*. We used Markov Chain Montecarlo (MCMC) to simulate the system evolution, introducing a temperature that, in our analogy, should set the “potency”.

We estimated the ID trend with temperature using a PCA-based estimator (Supplementary Eq. 6). The simulations were repeated 10 times with different random initialization, and we report in Supplementary Fig. 2A the mean values and the standard deviations (shaded area) of the estimated IDs. Despite the fluctuations, the average ID has precisely the expected trend: it decreases if we “freeze” the system. This is not simply due to the occupation of different basins of attraction (Supplementary section 2.2), but rather a consequence of the increased geometrical constraints. In fact, the trend is the same if we consider one single stored memory and we initialize all trajectories in this minimum, thus exploring a single basin of attraction at different temperatures (Supplementary Fig. 2B). Using the Hopfield model, we can also test how the ID estimation depends on the number of minima. As the number of attractors increases, PCA-based estimators are expected to report a significant decrease in the ID, as the directions between attractors capture a large portion of the variance. To test this, we ran simulations of the Hopfield model at a fixed temperature but with a variable number of stored patterns (ranging from 1 to 20, as shown in Supplementary Fig. 2C). This setup is designed to mimic the effect of increasing the number of cell types. We observed that the ID initially decreases with the number of attractors, which aligns with our intuition. As the number of memories increases further, the ID appears to slightly recover and stabilize. Ideally, with a very large number of attractors, the ID would be constrained only by the sample size.

It is important to note that, unlike in transcriptomic data in presence of multiple cell types, the heterogeneity among the basins of attraction in this model is likely much smaller. The different memories stored in the Hopfield model are equivalent random patterns, whereas transcriptomic profiles likely exhibit meaningful structure and substantial heterogeneity in the intrinsic dimension of the manifold corresponding to different cell types. These factors can easily explain the quantitative difference in the behavior of the ID with respect to the number of attractors in the model and in real transcriptomic data.

### 4 The relation between intrinsic dimension and gene-gene correlations

The observation that the ID decreases with differentiation is coherent with the Waddington landscape picture and with the idea that the expression profiles of differentiated and specialized cells are highly regulated. Gene regulation should, in turn, induce observable correlations between gene expression levels. Equivalently, from a landscape perspective, the presence of geometrical constraints should induce correlation between features. Indeed, the increase in heterogeneity of the eigenvalues of the covariance matrix, which we observe using PCA-based estimators during differentiation, also generally correspond to an increased level of feature covariance.

While the intrinsic dimension well recapitulates the global geometrical properties of the dataset, we can anyway analyze the gene-gene correlation structure during development as a consistency check. To this aim, we can define the network of genegene correlation and study its evolution. Specifically, for each developmental stage, we first selected the first 3000 highly variable genes - to reduce the computational complexityusing *scanpy* standard pre-processing function, setting *seurat v3* as flavor. We then constructed a gene-gene co-expression network by weighting each link with the Pearson correlation coefficient *ρ*. Links were pruned if |*ρ*| was smaller than a specified threshold, for example 0.4, 0.5 or 0.6. The number of non-isolated genes (genes with at least one link with a weight exceeding the threshold) can be used as a proxy for the global level of correlation. Supplementary Fig. 4 shows how this quantity increases during mouse gastrulation [34], suggesting a gradual increase in gene-gene correlation with development and supporting the idea that differentiation is accompanied by tighter gene regulation.

However, the number of sufficiently correlated gene pairs only captures a specific aspect of the data statistics during differentiation. For example, all genes are con-sidered equally important, regardless of their average expression level or variance. In contrast, the intrinsic dimension appears to be a more general and straightforward measure of the global geometrical properties of the data, which can be correlated with biological properties such as cell potency.

### 5 Intrinsic dimension versus alternative differentiation potential correlates

This section analyzes two previously proposed correlates of cell potency that, like intrinsic dimension, are based on simple statistical/geometrical properties of expression profiles without requiring prior biological knowledge.

A scRNA-seq experiment provides a transcript count matrix *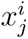*, with *i* = 1, 2*, …, N* (*N* is the number of cells) and *j* = 1, 2*, …, D* (*D* is the number of genes). *x^i^* is the vector representing the expression profile of the i-th cell. Gulati et al. [66] observed that the number of detected genes (i.e., the genes with at least one detected transcript) decreases with cell differentiation (Figure 1 of ref. [66]). They called this quantity Transcriptional Diversity (TD), and it is defined for every cell *i* as *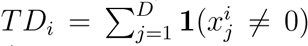*, i.e. the number of counts in the cell that are not zero. A possible rationale behind the TD trend with potency is that stem cell are less regulated and thus have more diverse expression profiles with respect to strongly regulated differentiated cells.

Another quantity, derived from statistical physics, that have been shown to correlate with cell potency in some datasets is the entropy of expression profiles [60, 61]. The entropy of single-cell expression profiles is defined as *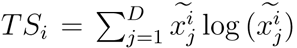*, total number of reads assigned to the cell, i.e. *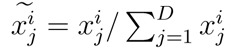*.

Transcriptional diversity and expression entropy form the basis of much more sophisticated tools, such as CytoTRACE [66] and StemID [61], which also incorporate additional information into their pipelines. However, our aim is not to benchmark these tools in detail, but rather to analyze and test the easily computable (and interpretable) statistical properties on which they are built as proxies of cell potency, and to compare them with intrinsic dimension.

While these two quantities (transcriptional diversity and entropy) sometimes correlate with cell potency in the dataset we analyzed, the correlation is less systematic compared to the ID-score. Supplementary Fig. 9 shows two examples of comparisons between ID-score, TD and TS in recovering differentiation potentials in mouse gastrulation [34] and pancreatic endocrinogenesis [41]. While the ID-score can reproduce the known hierarchies, in these examples both TD and TS produce incorrect rankings. Sometimes the expected hierarchy is even reversed.

It is important to notice that both the transcriptional diversity and the expression entropy are properties of single cells, while the ID-score is a score for a group of cells. In fact, Supplementary Fig. 9 compares the distributions of TD and TS as box plots, with the ID measured over different sub-samplings. This difference can make the ID a more robust measure of global properties of the dataset, less sensitive to noise from stochastic gene expression and from technical origins. The downside is that the ID-score cannot attribute a potency score to every single cell, but requires a preliminary cell grouping or clustering.

Moreover, entropy and transcriptional diversity are strongly dependent on number of detected genes. Transcriptional diversity is directly defined by this quantity, while the maximum value of entropy depends on this quantity. However, the number of detected genes is highly affected by the sequencing depth and by the sampling process involved in RNA sequencing. A global geometric measure, such as the ID, should be less sensitive to data sparsity due to sampling. These two factors could explain the higher robustness of the ID-score as a proxy for potency that we observed in the analyzed datasets.

### 6 Description of the analyzed datasets

The freely-available scRNA-seq datasets that we considered for our analyses are largely from the Gene Expression Omnibus repository and rely on experimental protocols using Unique Molecular Identifiers (UMIs). We considered datasets, listed below, that refers to both whole embryos or single organs or tissues as a function of time. In most of the datasets, cells are annotated by the authors depending on both the developmental stage and the cell type.

#### Mouse Cell Atlas

Mouse cell atlas [37, 38] is a dataset obtained via Microwell-seq. Hundreds of single cells have been sequenced covering all of the major mouse organs and the cell types have been identified using a clustering procedure. The analyzed time-stages are: fetal (embryonic day E14.5), neonatal, ten days, three weeks, and adult (6-10 weeks).

We considered the 8498 protein-coding genes that have been sampled in every cell of every organ.

#### Mouse Haematopoietic stem/progenitor cells

In the homeostasis of several adult tissues, multipotent progenitor cells continuously differentiate into specialized cell types in a continuous self-regulating process of regeneration and renewal. For example, all cellular blood components are derived from hematopoietic stem cells during the hematopoiesis.

In [46], 44.802 single-cell transcriptomes are reported, covering the hematopoietic stem/progenitor (HSPC) compartment from mouse bone marrow. The sequencing protocols are Smart-Seq2 and 10x Chromium.

To obtain a comprehensive view of hematopoiesis, the cells are sorted in two broad gates: LK, capturing HSPCs, and Lin^-^Sca-1^+^c-Kit^+^ (LSK), a subset of the LK gate enriched for more immature progenitors. The sorting gate to isolate LSK and LK cells was based on c-Kit and Sca-1 surface expression for droplet-based scRNA-seq.

#### Mouse Gastrulation and early organogenesis

Ref. [34] investigates the dynamic process of cellular diversification during gastrulation and early organogenesis of mice that occurs in the 48 hours spanning from embryonic day (E) 6.5 to E8.5. Profiles from whole mouse embryos were collected at six-hour intervals between E6.5 to E8.5. In total, 116.312 single-cell transcriptomic profiles were clustered and annotated, identifying 37 major cell populations. ScRNA-seq libraries were generated using the 10x Genomics Chromium system (v.1 chemistry) and samples were sequenced on an Illumina Hi-Seq 2500 platform.

In this dataset, the information about the specific biological samples is provided. Therefore, to reconstruct the temporal ID trend of Fig. 1B, we restricted our analysis to cells coming from the same sample (for each developmental stage), in order to reduce possible batch effects. We could not do the same distinction for Fig. 3B due to the limited number of cells.

#### Mouse pancreatic endocrinogenesis

The formation of pancreatic endocrine cells occurs in two distinct stages during rodent embryonic development. The first stage occurs between E9.0 and E12.5, while the second one between E12.5 and E15.5, during which the majority of endocrine cell types are formed. With the aim of resolving the molecular changes during endocrinogenesis, high-throughput scRNA-seq analysis of 38.000 pancreatic epithelial cells during the secondary transition (E12.5, E13.5, E14.5, and E15.5) was performed [41]. More specifically, four scRNA-seq experiments were performed using 10X Genomics technology. Unsupervised graph-based clustering revealed eight major cell clusters including multipotent pancreatic progenitors (MPCs), tip, trunk, acinar, ductal, EPs, Fev^high^, and endocrine cells. Cell clusters were annotated using the expression of well-known marker genes and the lineage relationships between these cell types were fully reconstructed [41].

#### Mouse corticogenesis

To build a comprehensive single-cell transcriptional and epigenetic atlas of the developing mammalian cerebral cortex, the authors of [42] performed scRNA-seq experiments over the entire period of corticogenesis: E10.5 and E11.5 (symmetrically dividing neuroepithelial cells); E12.5 and E13.5 (birth date of layer 6 and 5 excitatory neurons); E14.5 to E17.5 (birth date of layer 4 and 2/3 excitatory neurons); and E18.5, postnatal day (P) 1 and P4 (gliogenesis). They report 98.047 scRNA-seq profiles, which include all known cell types of the developing cerebral cortex.

They used a pseudotime approach based on diffusion (URD) [36], to generate a branching tree of trajectories based on the transcriptional similarity of pseudotime ordered cells. The root consists of early progenitors belonging to E10.5, the tips come from P4 and correspond to terminal cell types.

Here, we focus on the embryonic time stages of corticogenesis, from E10.5 to E18.5. We also investigate how the ID-score recapitulates the differentiation trajectories of excitatory neurons in the two progressive stages: from apical to intermediate progenitors, until the state of excitatory neurons.

#### Mouse embryoids

In Ref. [40], they applied tiny sci-RNA-seq3 to profile 285.640 single cell transcriptomes of multiple individual ‘ETiX’ mouse embryoids assembled from embryonic stem cells, trophoblast stem cells and inducible extraembryonic endoderm stem cells. The data concern the day 6 and day 8 of development. Amadei et al. also studied scRNA-seq data of natural mouse embryos at E7.5, E8, E8.5, E8.75. and E9. The aim of their work consisted in showing that ETiX mouse embryoids can firstly develop into gastrulating embryoids, and secondly into neurulating embryoids. In conclusion, they found out that the ETiX embryoids recapitulate the development of whole natural mouse embryos in uterus up to day 8.5 post-fertilization.

In our study, we only used data from embryoids to compare the value of ID at day 6 and 8 (Fig. 2E).

#### Zebrafish embryogenesis - Wagner

In the study [33], over 92.000 cells from zebrafish embryos were subjected to inDrops single-cell RNA sequencing throughout the initial day of development. Employing a graph-based methodology together with analysis of known marker genes, the researchers delineated a cell-state landscape elucidating axis patterning, germ layer formation, and organogenesis. The data come from 7 different developmental stages (4 hours post-fertilization, 6 hpf, 8 hpf, 10 hpf, 14 hpf, 18 hpf and 24 hpf).

#### Zebrafish embryogenesis - Farrell

In [36], Farrell et al. studied single-cell transcriptomes of 38.731 cells obtained with Drop-seq during early zebrafish embryogenesis at a high temporal resolution, spanning 12 stages from the onset of zygotic transcription through early somitogenesis. To identify the transcriptional trajectories in the data they developed a simulated diffusion-based computational approach (URD), which identified the trajectories describing the specification of 25 cell types in the form of a branching tree, where the the root of the tree correspond to a multipotent cell type, while terminally differentiated cell types constitutes the tips.

#### Zebrafish neurogenesis

A 2020 study by Raj et al. [43] analyzed the gene expression of over 220,000 individual cells isolated in zebrafish brains, using the 10X Chromium scRNA-seq platform. These cells originated from 12 distinct developmental stages, spanning from embryo to larva. Analyses of known and novel marker genes revealed about 800 clusters and allowed to characterize the transition from progenitors to neurons and, more in general, the molecular mechanisms underlying vertebrate neurogenesis.

#### Zebrafish embryogenesis - Farnsworth

In [44], the study was conducted on 44.102 cells coming from 6 different experiments and concerning the development of zebrafish embryos. From single-cell RNA sequencing data, Farnsworth et al. computationally reconstructed 220 clusters and annotated them by analysing the expression of some marker genes, identified through ZFIN database (Zebrafish Information Network). ScRNA-seq libraries were generated using the 10x Genomics Chromium platform (v.2 chemistry) and samples were sequenced on either an Illumina Hi-Seq or an Illumina Next-seq.

In Fig. 3D, we compare the ID of three classes of cell types (being represented at least by 180 cells) that take part in the genesis of retinal neurons and photoreceptors. These classes are retinal progenitors (RetProgAlla, RetProg1a, RetProg0a, RetProg0b, RetProgAllb, RetProgAllc), differentiating retinal cells (RetDiff2, RetDiff25a, RetDiffAll, RetDiff25b, RetDiff25c, RetDiff25d, RetDiff25e) and retinal neurons (RetNeuron25, RetPR - retinal photoreceptors).

#### Hydra turnover

The cnidarian polyp Hydra undergoes continual self-renewal and is capable of whole-body regeneration from a small piece of tissue. The stem cell populations, morphological cell types and lineage relationships are well characterized. In [45], they sequenced 24.985 single-cell transcriptomes and identified the molecular signatures of a broad spectrum of cell states, from stem cells and progenitors to terminally differentiated cells, building differentiation trajectories for all cell lineages. In our work, for the two epithelial layers (endoderm and ectoderm) and the interstitial lineage we compared the ID of progenitors and differentiated cell clusters, selecting only sufficiently represented cell types having more than 330 cells.

In Fig. 4A, the points refer to cell types belonging to the interstitial layer and are ordered in the following way: stem/progenitor cells, neuronal cells (nc) progenitors, nc and gland cells (gc) progenitors, differentiated nematocyte, ectodermal neuron (n ec) 1, n ec 2, male germline, granular mucous gland cells, spumous mucous gland cells 2, zymogen gland cells 1. Moving on to endodermal epithelilal cells, in Fig. 4B, points are respectively referred to: stem cells (SC) 1, SC2, SC3, cells from head, cells from foot, cells from tentacles. Finally, the order chosen for ectoderm in Fig. 4C is: SC1, SC2, differentiated nematocyte, cells from head, basal disk, multiplets of battery cells 2, nematoblasts (suspected phagocytosis doublet).

#### C. elegans embryogenesis

In [35], the authors sequenced the transcriptomes of single cells from C. elegans embryos with the 10X Genomics platform. They assayed loosely synchronized embryos enriched for preterminal cells as well as embryos that had been allowed to develop for ∼300, ∼400, and ∼500 min after the first cleavage of the fertilized egg. After the quality control, they estimated the embryo stage of the 86.024 single cells by comparing their expression profile with a high-resolution whole-embryo RNA-seq time series [79].

### 7 Supplementary Table

**Supplementary Tab. 1:** Description of the datasets. The number of cells sub-sampled from each group (p.s.=per stage, p.ct.=per cell type, p.b.s.=per biological sample, p.u.s=per URD segment) is specified for every analyzed dataset (Figs. 1 2 3 4). The total number of detected genes and protein-coding genes (defined by the Ensembl database [71]) is reported. In the last two columns, we indicate the two parameters (Eq. (2) and (3)) used to define the normalized ID-score.

| Dataset | Number of cells |  | Number of genes |  | Normalization |  |
| --- | --- | --- | --- | --- | --- | --- |
| | Total | sub-sampled | Total | Protein-coding | $Y_{\text{shift}}$ | $Y_{\text{scaling}}$ |
| Mouse gastrulation [34] | 108839 | 1530 p.s.<br>467 p.ct. | 29452 | 23727 | 34<br>14 | 60<br>46 |
| Mouse Pancreas [41] | 36351 | 3781 p.s.<br>918 p.ct. | 27998 | 16558 | 28<br>18 | 50<br>49 |
| Mouse cell atlas [37, 38] | 231732 | 5000 p.s. (organism)<br>1478 p.s. per organ | 9348 | 8498 | 40<br>36 | 70<br>84 |
| Mouse corticogenesis [42] | 77842 | 781 p.s.<br>4995 p.ct. | 27998 | 16461 | 31<br>39 | 106<br>60 |
| Mouse hematopoiesis [46] | 44802 | 5000 p.b.s. | 27998 | 16109 | 59 | 78 |
| Mouse embryoids [40] | 285640 | 1464 p.b.s. | 49585 | 17847 | 54 | 114 |
| Zebrafish embryogenesis [36] | 38279 | 142 p.s.<br>154 p.u.s. | 23974 | 14081 | 27<br>18 | 77<br>58 |
| Zebrafish embryogenesis [44] | 44020 | 136 p.ct. | 32520 | 23911 | 9 | 46 |
| Zebrafish embryogenesis [33] | 36749 | 2676 p.s. | 30677 | 14059 | 37 | 87 |
| Zebrafish neurogenesis [43] | 137706 | 5000 p.s. | 32191 | 22905 | 22 | 60 |
| Elegans embryogenesis [35] | 86024 | 344 p.s. | 20222 | 18000 | 8 | 27 |
| Hydra turnover [45] | 24984 | 332 p.ct. interstitial<br>344 p.ct. endoderm<br>337 p.ct. ectoderm | 36814 | NA | 8<br>16<br>15 | 48<br>32<br>26 |

### 8 Supplementary Figures

#### 8.1 Supplementary Fig.1

**Supplementary Fig. 1:**
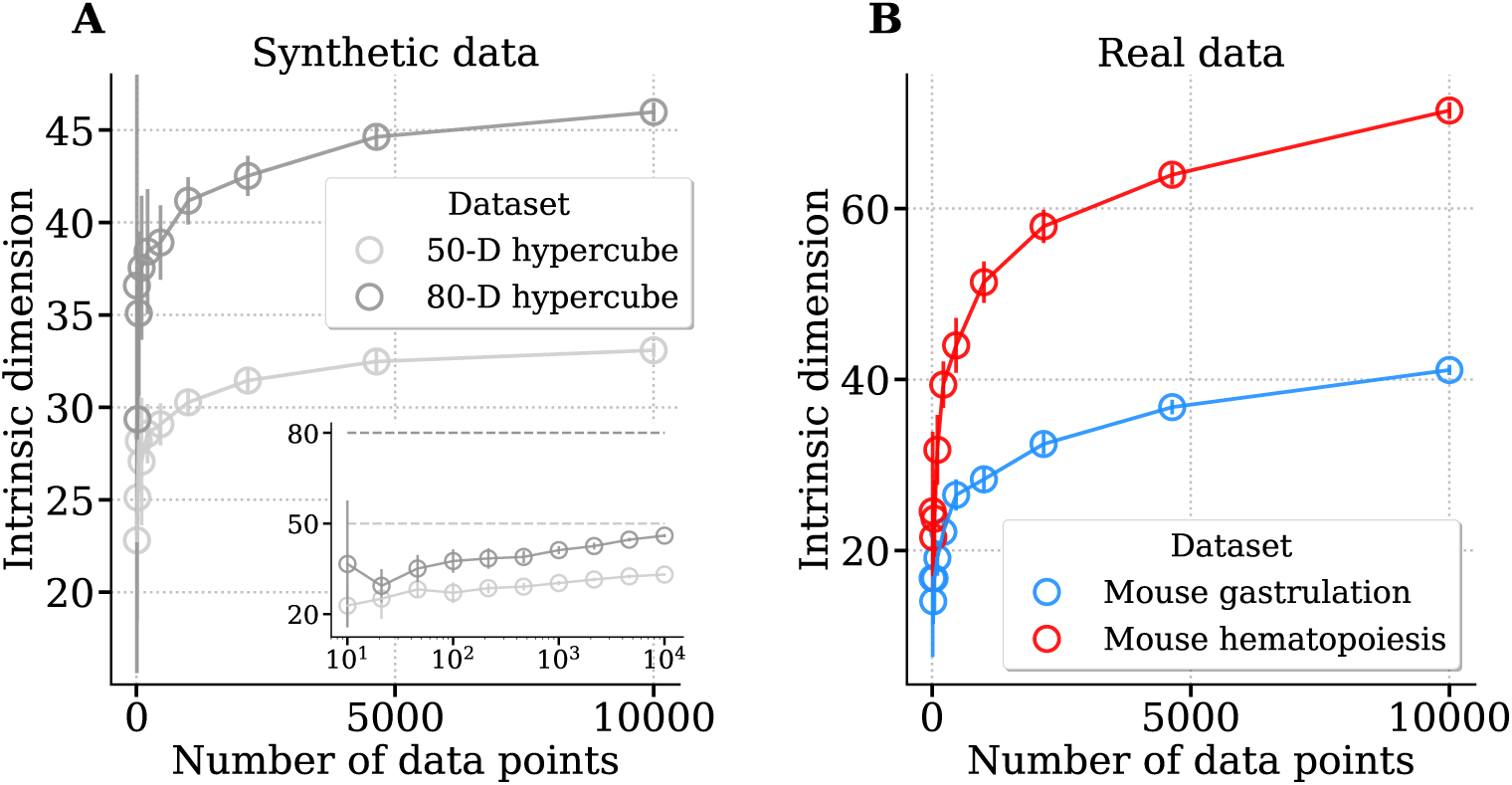
Relationship between the estimated ID and the sample size. **A**) 50-dimensional and a 80-dimensional hypercubes embedded in a 1000-dimensional space are considered. A variable number of points is sampled from these two synthetic datasets. The ID estimations can correctly rank the two manifold dimensions if evaluated at the same sample size. However, the inset shows that the estimated values are far from the true ID (dotted line). **B**) The same analysis is repeated on two real datasets related to mouse gastrulation [34] and mouse hematopoiesis [46] dataset.

#### 8.2 Supplementary Fig.2

**Supplementary Fig. 2:**
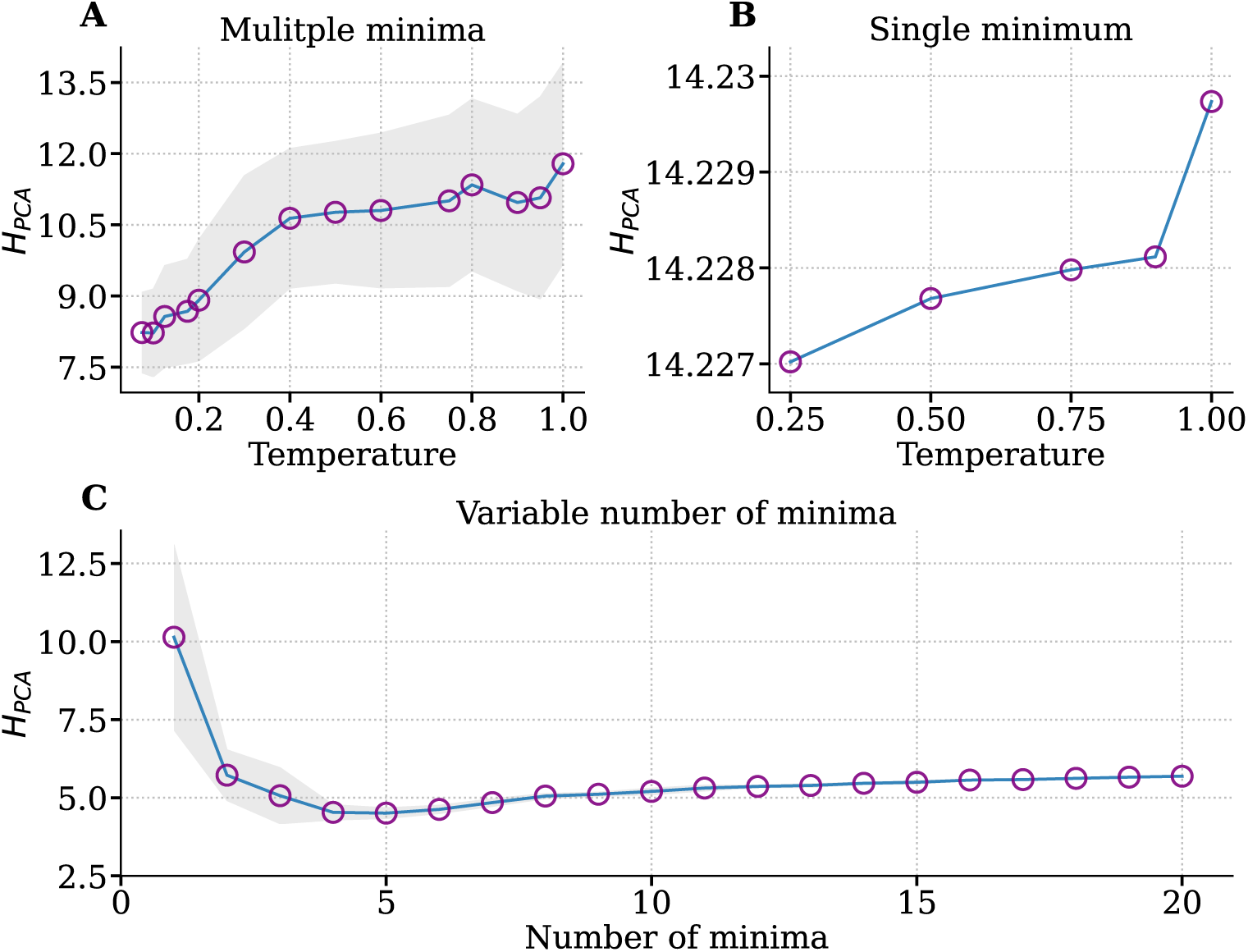
Intrinsic dimension dependence on temperature and on the number of memories in the Hopfield model. **A**) ID values are obtained with the *H_PCA_* estimator (Eq. 6). We simulated a fixed number of 1500 randomly initialized trajectories of the Hopfield model with 10 random memory stored at different temperature values. We computed the mean *H_PCA_* values over 10 independent simulations and considered a 3-points-window moving average, while the shaded area reports the standard deviations. As hypothesized, the intrinsic dimension decreases as the temperature decreases. **B**) The same simulations are performed with a single random memory stored and with all trajectories initialized in this minimum. Shaded area is not visible because error bars on y values have order magnitude 10^−15^. **C**) With a fixed temperature of *T* = 0.1, the *H_PCA_* is calculated as a function of the number of attractors (memories stored) in the energetic landscape, from 1 to 20. In this case, we consider 250 trajectories and mean values are computed over 10 simulations.

#### 8.3 Supplementary Fig.3

**Supplementary Fig. 3:**
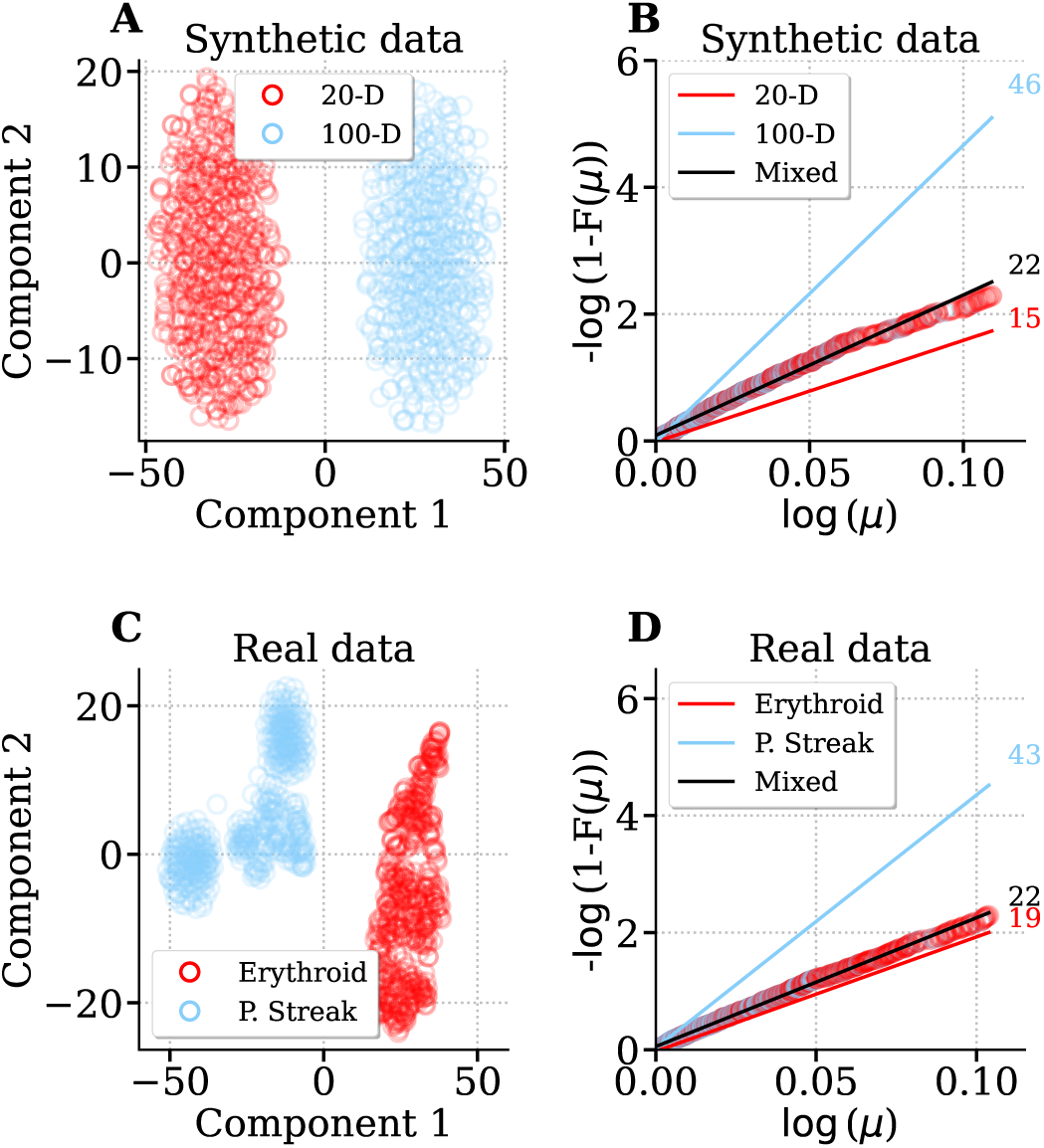
In the presence of multiple data manifolds ID estimators are biased towards low dimensions. In the first row: data sampled from a 20 and 100-dimensional hypercube embedded in a 1000-dimensional space, each one represented by 10^3^ points. In the second row: data from two cell types (erythroid and primitive streak) from the mouse gastrulation dataset [34], with 10^3^ cells per cell type. **A** and **C**) The data projection on the 2D space given by t-SNE shows the presence of two well separated clusters. **B** and **D**) Relationship between *µ* (introduced in Eq.(4)) and its cumulative distribution *F* (*µ*). The pale blue line and the red one are the fitted lines when single manifolds are considered. The black line is obtained taking the whole dataset. The values reported on the right correspond to the ID estimations based on TWO-NN. Note that we are in the under-sampled regime.

#### 8.4 Supplementary Fig.4

**Supplementary Fig. 4:**
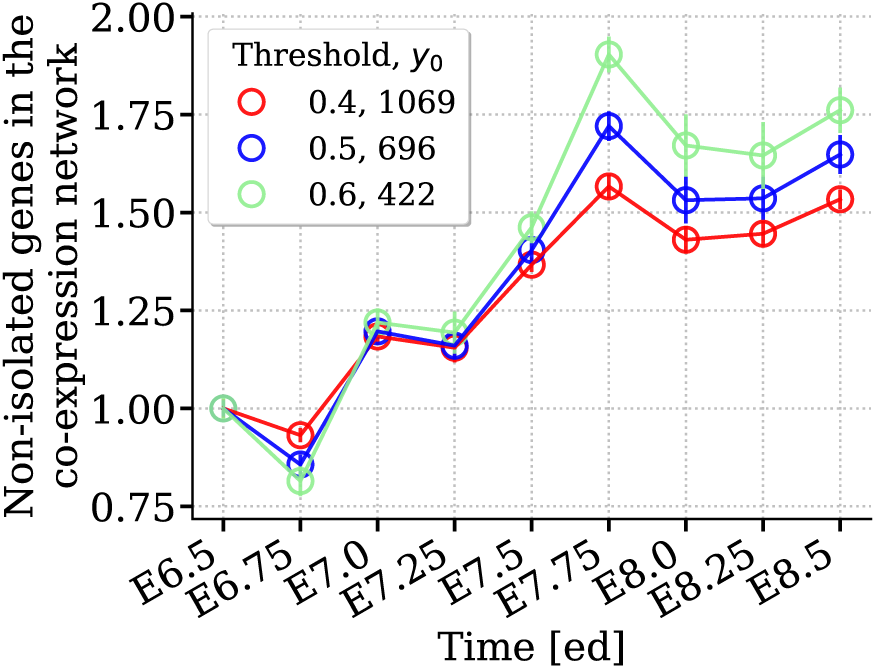
The correlations among gene expression levels increase during development. The number of genes in the co-expression network grows with developmental stages in the mouse gastrulation dataset [34]. We report the trend for different threshold values used to construct the network. Values on the y-axis are scaled by the value corresponding to stage E6.5 (*y*_0_), as indicated in the legend.

#### 8.5 Supplementary Fig.5

**Supplementary Fig. 5:**
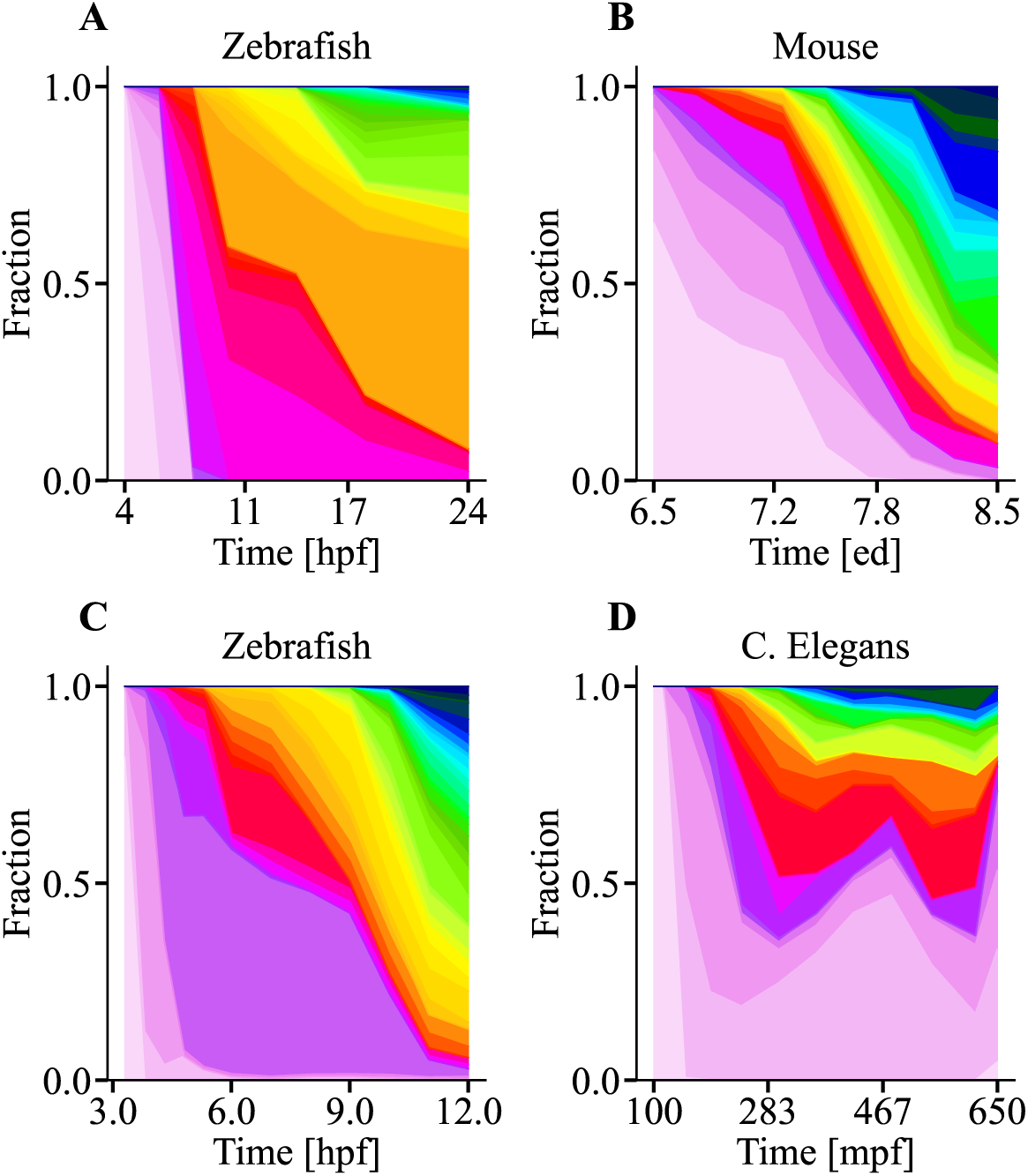
The number of cell types increases during embryogenesis. Cell-type proportions during pre-natal development for Zebrafish 4-24 hpf [33] (**A**), Mouse 6.5-8.5 ed [34] (**B**), Zebrafish 3.3-12 hpf [36] (**C**), and C. Elegans 100-650 mpf [35] (**D**). For each animal, each color corresponds to a specific cell type as annotated by the authors. The number of distinct colors increases as a function of developmental time.

#### 8.6 Supplementary Fig.6

**Supplementary Fig. 6:**
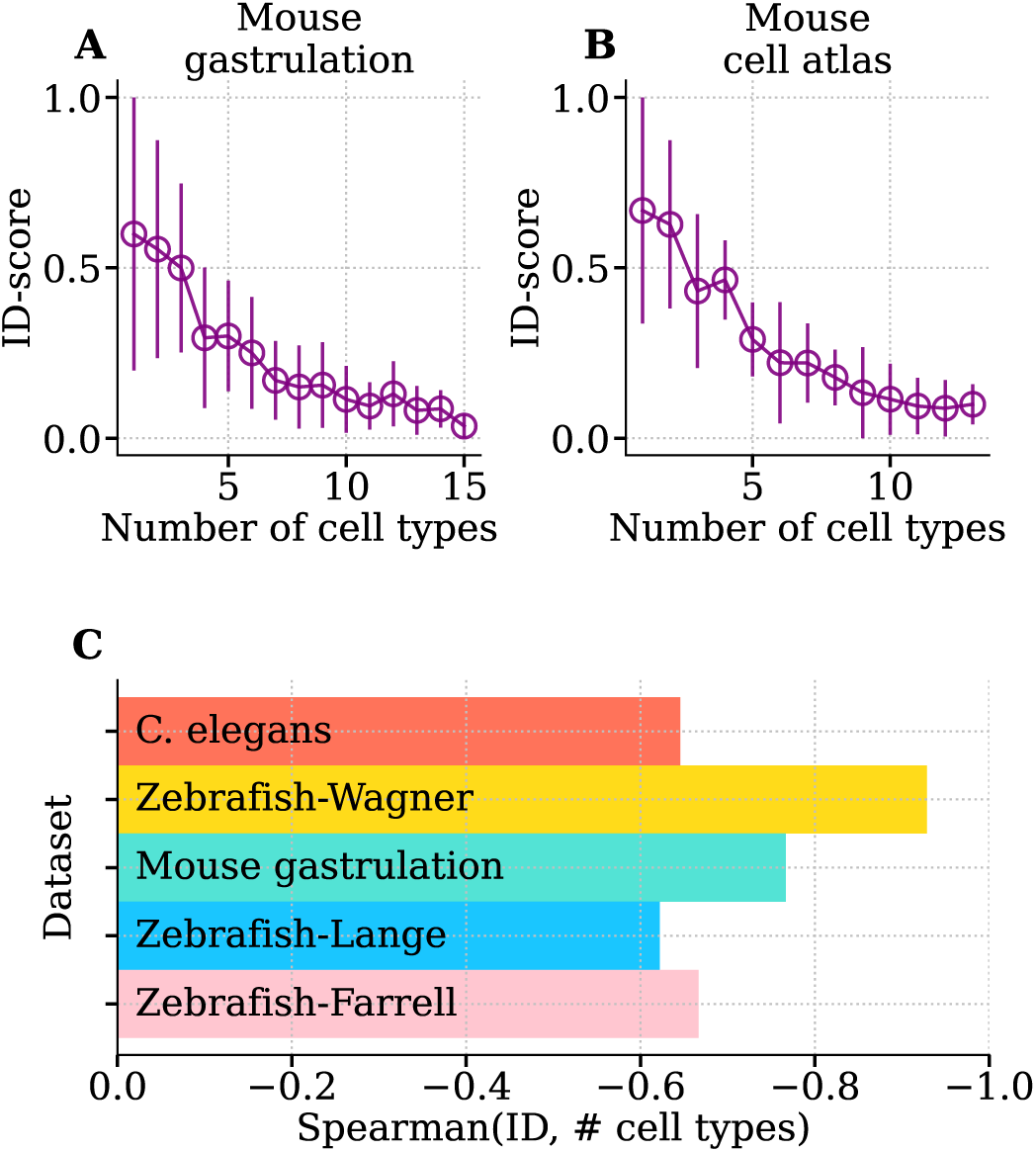
The ID decreases with the number of cell types independently of their overall level of differentiation. We report the ID-score (symbol ± error bar = mean ± standard deviation) of controlled mixtures of cell types, obtained by randomly assembling cell types from the Mouse cell atlas [37, 38] (**A**) and the Mouse gastrulation dataset [34]. **C**) Spearman correlation between the number of cell types and the ID-score calculated for different datasets relative to embryonic development.

#### 8.7 Supplementary Fig.7

**Supplementary Fig. 7:**
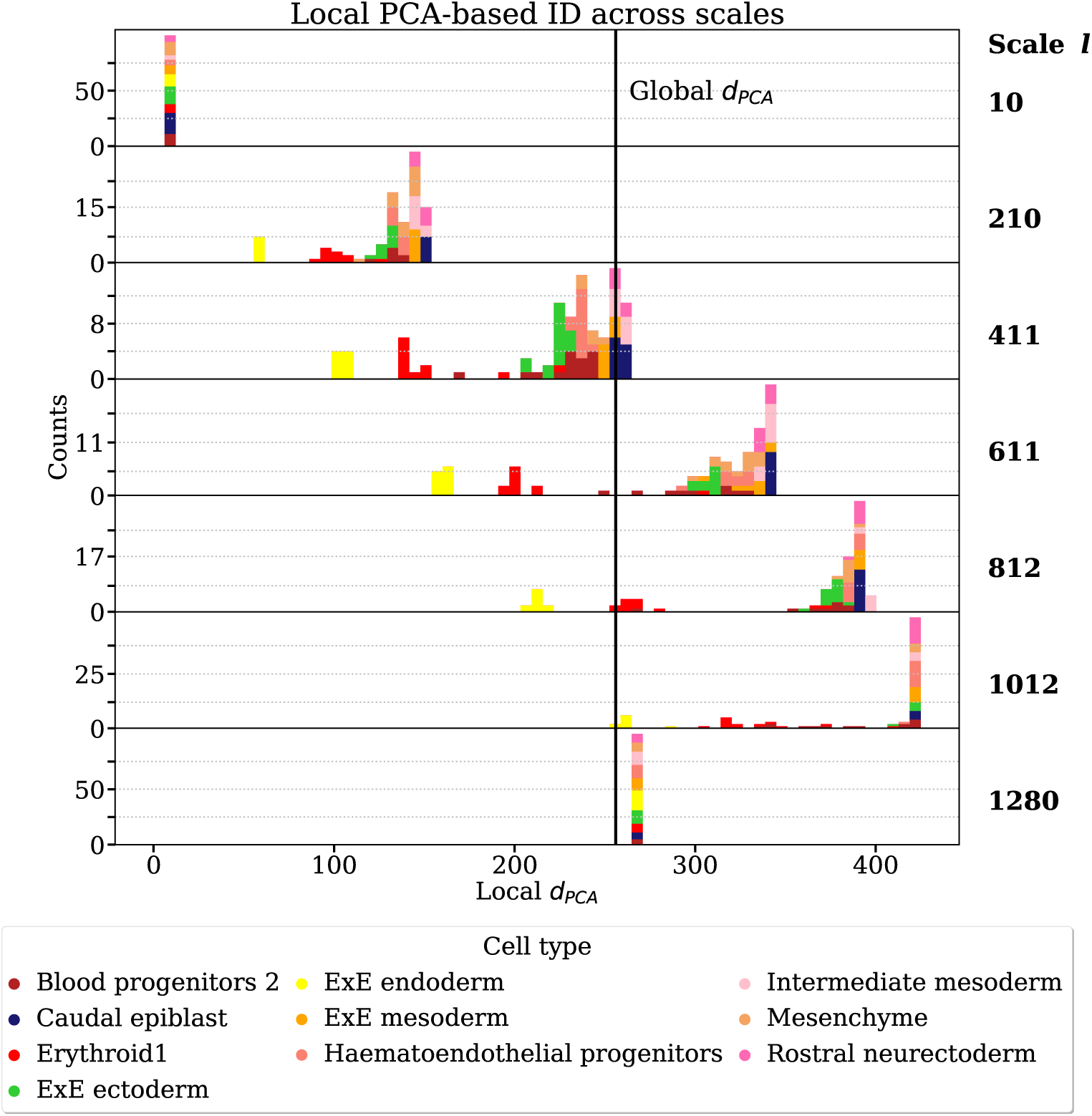
**A local PCA-based estimator reveals cell types heterogeneity across different scales**. The local version of *d_PCA_*, i.e., *Ld_PCA.l_* is applied at different scales (*l*) on 1300 cells from embryonic day 8 of mouse gastrulation [34]. at each scale the scale, we measured the local ID on 50 sub-samples of randomly selected cells. The colours encode for the cell type. Each cell type is equally represented by 130 cells. The vertical black line specifies the value of *d_PCA_*taking the 1300 cells together.

#### 8.8 Supplementary Fig.8

**Supplementary Fig. 8:**
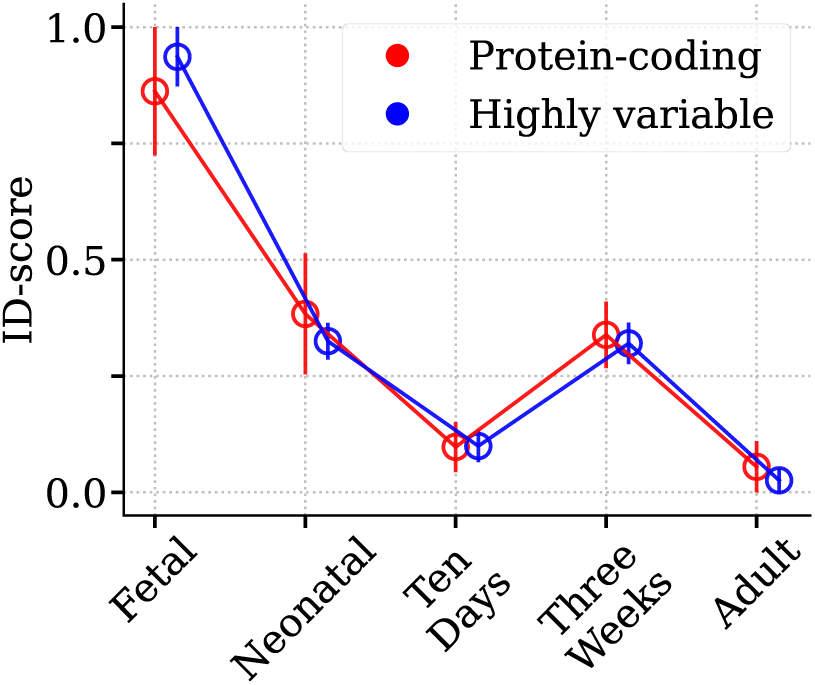
Robustness of results with respect to feature selection. ID measured for different temporal stages of the Mouse Cell Atlas dataset [37, 38]. We firstly took into account all 8498 protein-coding genes (red curve), then we restricted the analysis to the first 2000 highly-variable genes detected with *seurat v3* of *scanpy* (blue curve).

#### 8.9 Supplementary Fig.9

**Supplementary Fig. 9:**
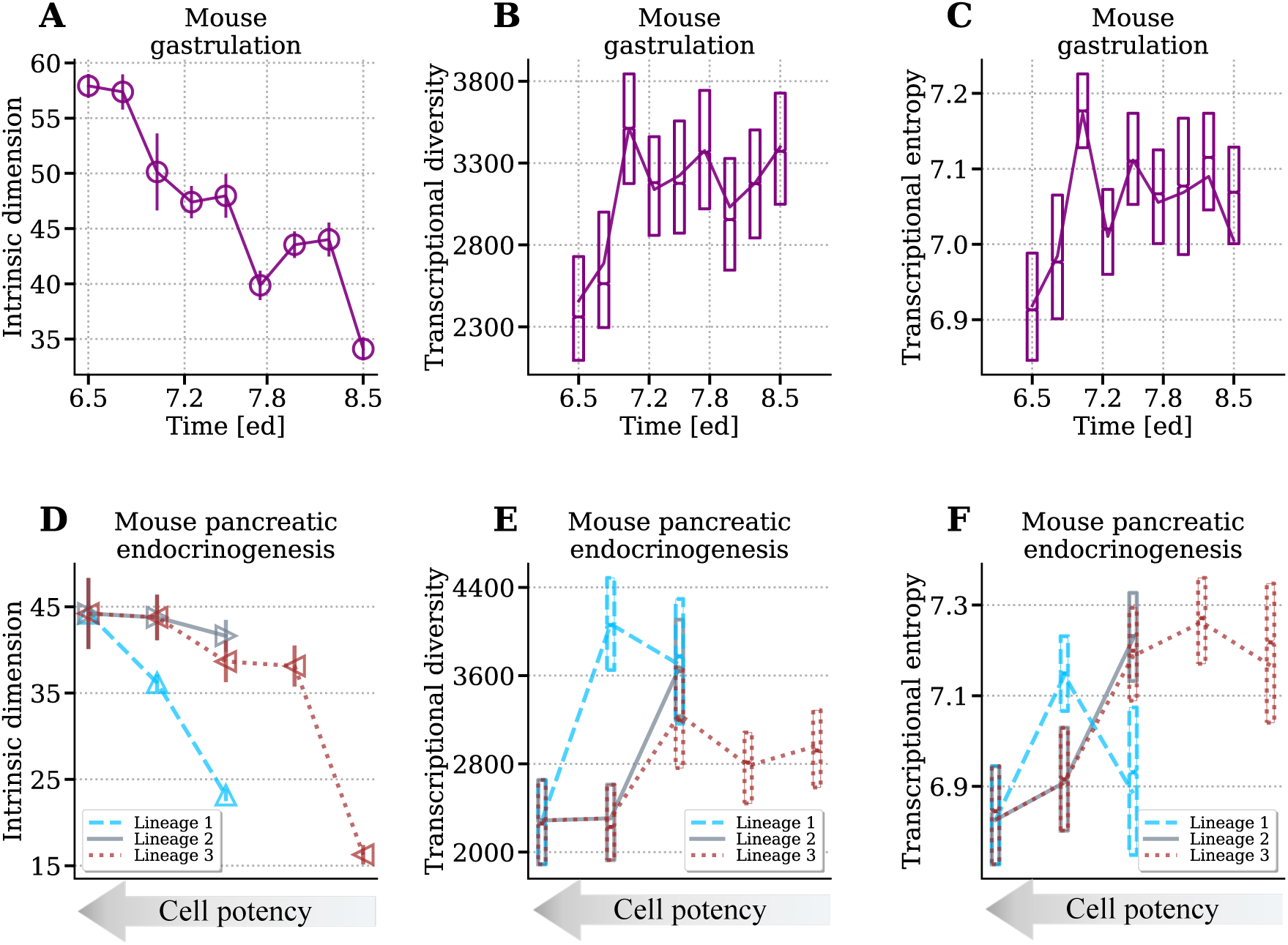
Comparison between ID, transcriptional diversity and transcriptional entropy. First row: cells from mouse gastrulation [34] grouped per developmental stage as in Fig. 1B. Second row: the hierarchy of cell types induced by the known differentiation diagram of pancreatic endocrinogenesis [41] (Fig. 3A), composed by lineage 1 (Multipotent-Tip-Acinar), lineage 2 (Multipotent-Trunk-Ductal) and lineage 3 (Multipotent-Trunk-Ep-Fev+Endocrine). Different lines connect the three lineages as in Fig 3. For each dataset, ID trends are shown (Figures **A** and **D**), together with boxplots resuming transcriptional diversity and transcriptional entropy distributions (Figures **B**,**E** and **C**,**F**).

#### 8.10 Supplementary Fig.10

**Supplementary Fig. 10:**
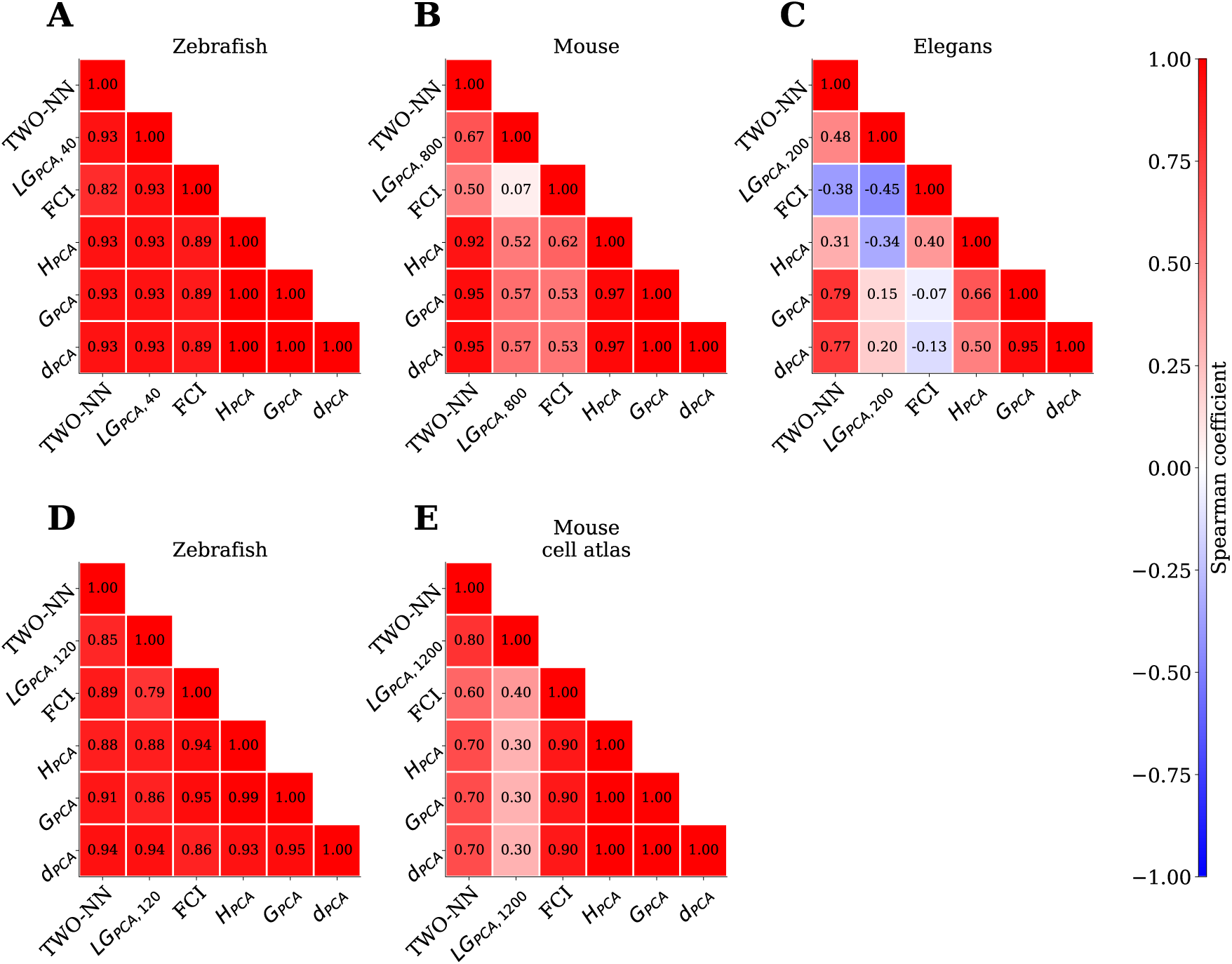
Correlation between temporal ID trends in whole organisms obtained with different ID estimators. Each trend showed in Fig. 1 is reproduced with a different ID estimator and the Spearman correlation coefficient is computed for every pair of estimators.

#### 8.11 Supplementary Fig.11

**Supplementary Fig. 11:**
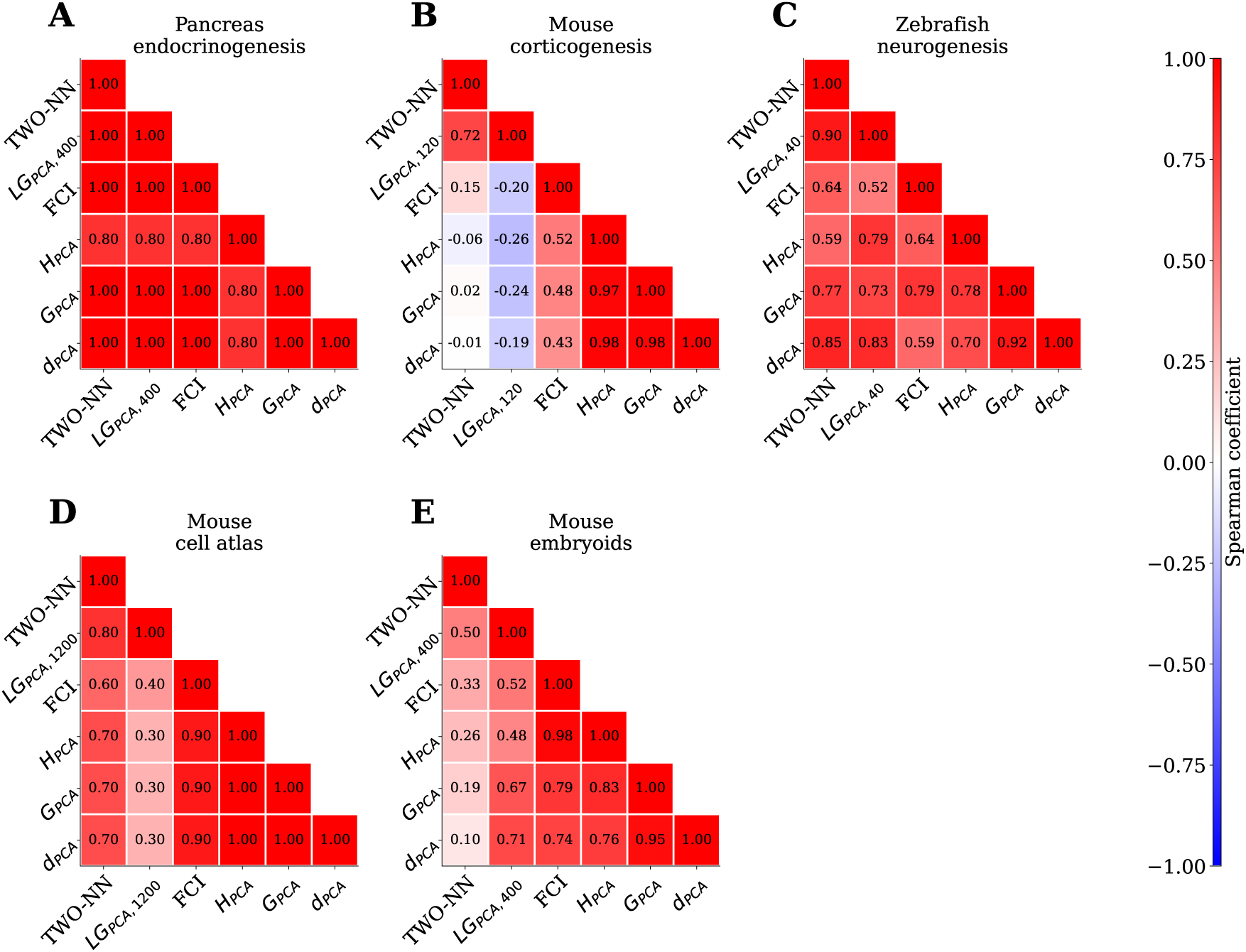
Correlation between temporal ID trends in single organs obtained with different ID estimators. Each trend showed in Fig. 2 is reproduced with a different ID estimator and the Spearman correlation coefficient is computed for every pair of estimators.

#### 8.12 Supplementary Fig.12

**Supplementary Fig. 12:**
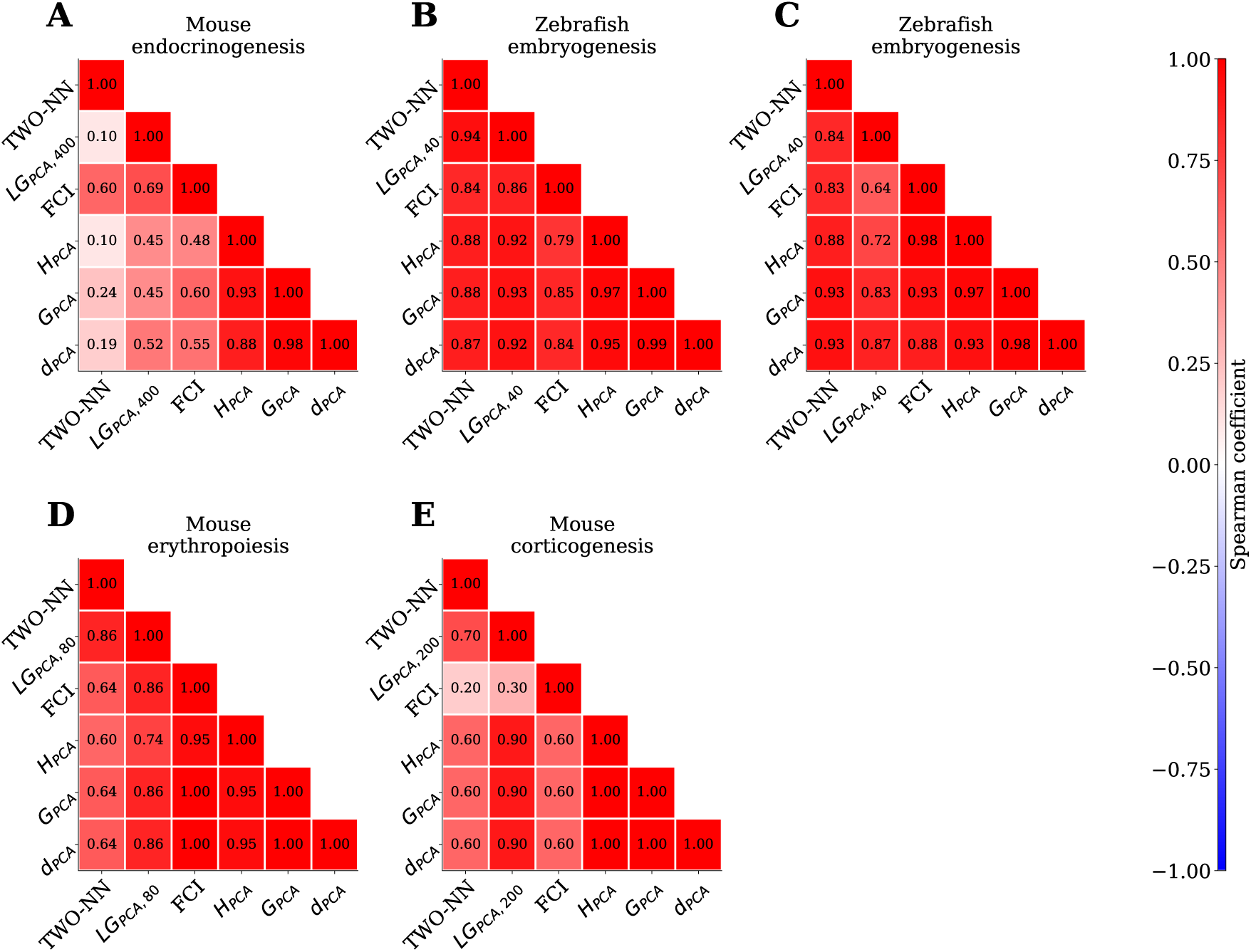
Correlation between ID values of cell types obtained with different estimators. Each trend showed in Fig. 3 is reproduced with a different ID estimator and the Spearman correlation coefficient is computed for every pair of estimators.

#### 8.13 Supplementary Fig.13

**Supplementary Fig. 13:**
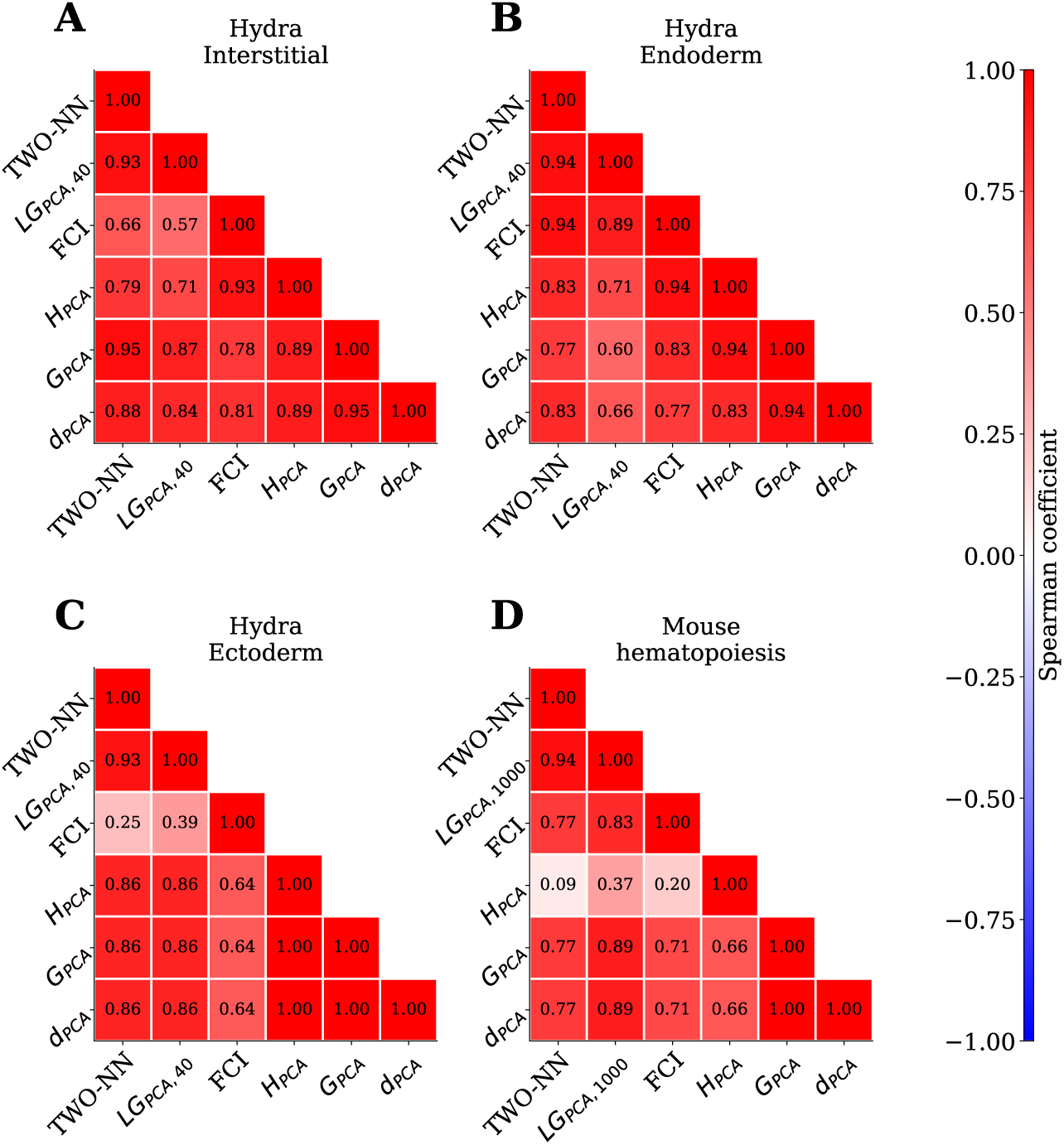
Correlation between ID values of cell types during tissue renewal obtained with different estimators. Each trend showed in Fig. 4 is reproduced with a different ID estimator and the Spearman correlation coefficient is computed for every pair of estimators.

